# Activation of the cGAS-STING innate immune response in cells with deficient mitochondrial topoisomerase TOP1MT

**DOI:** 10.1101/2022.03.08.483326

**Authors:** Iman Al Khatib, Jingti Deng, Yuanjiu Lei, Sylvia Torres-Odio, Gladys R. Rojas, Laura E. Newman, Brian K. Chung, Andrew Symes, Hongliang Zhang, Shar-yin N. Huang, Yves Pommier, Aneal Khan, Gerald S. Shadel, A. Phillip West, William T. Gibson, Timothy E. Shutt

## Abstract

The recognition that cytosolic mtDNA activates cGAS-STING innate immune signaling has unlocked novel disease mechanisms. Here, an uncharacterized variant predicted to affect TOP1MT function, P193L, was discovered in a family with multiple early-onset autoimmune diseases, including Systemic Lupus Erythematosus (SLE). Although there was no previous genetic association between *TOP1MT* and autoimmune disease, the role of TOP1MT as a regulator of mtDNA led us to investigate whether TOP1MT could mediate the release of mtDNA to the cytosol, where it could then activate the cGAS-STING innate immune pathway known to be activated in SLE and other autoimmune diseases. Through analysis of cells with reduced TOP1MT expression, we show that loss of TOP1MT results in release of mtDNA to the cytosol, which activates the cGAS-STING pathway. We also characterized the P193L variant for its ability to rescue several TOP1MT functions when expressed in *TOP1MT* knockout cells. We show that the P193L variant is not fully functional, as its re-expression at high levels was unable to rescue mitochondrial respiration deficits, and only showed partial rescue for other functions, including repletion of mtDNA replication following depletion, nucleoid size, steady state mtDNA transcripts levels, and mitochondrial morphology. Additionally, expression of P193L at endogenous levels was unable to rescue mtDNA release-mediated cGAS-STING signaling. Overall, we report a link between TOP1MT and mtDNA release leading to cGAS-STING activation. Moreover, we show that the P193L variant has partial loss of function that may contribute to autoimmune disease susceptibility via cGAS-STING mediated activation of the innate immune system.

## Introduction

The innate immune response is the first line of defense against various microorganisms and pathogens. However, the immune system sometimes recognizes intracellular components as foreign and attacks its own tissue and organs, leading to autoimmune diseases. The exact cause for such erroneous autoimmune responses is not always understood, as there are many triggers that can lead to the self-reactivity of the immune system. Systemic lupus erythematosus (SLE) is a chronic, heterogenous, systemic autoimmune disease, in which accumulating evidence suggests that mitochondrial dysfunction plays an important role(1, 2). Multiple genetic variants, including highly penetrant rare variants, have been associated with lupus susceptibility. The identification and functional characterization of such rare variants can provide significant insight into the cause, mechanisms, and potential treatment options for patients(3–6).

Mitochondria, best known as the powerhouses of the cell, are eukaryotic organelles that evolved from prokaryotes through an endosymbiotic event that occurred billions of years ago(7). The strongest evidence for their bacterial origin is the fact that mitochondria have retained their own 16,569 bp circular genome, the mitochondrial DNA (mtDNA), which is present in hundreds to thousands of copies in the cell(8). Release of mtDNA into the cytosol, in response to pathogen-induced mitochondrial stress, is a mechanism by which cells can amplify innate immune signaling in response to pathogen invasion(9, 10). However, in the absence of pathogens, mitochondrial dysfunction can also cause release of mtDNA into the cytosol, leading to ‘sterile’ activation of the innate immune system(11–13). The mechanism of release of mtDNA into the cytosol has been studied primarily in the context of apoptosis through BAK/BAX(14). However, another mechanism of mtDNA release has been described that depends on the opening of mitochondrial permeability transition (MPT) pores in the inner mitochondrial membrane(15), and oligomerization of voltage-dependent anion channel (VDAC) in the mitochondrial outer membrane(16). While less is known about when and how mtDNA is released under more physiological conditions, there is a growing appreciation of the functional consequences with respect to inflammation and disease. Recent research has demonstrated that release of mtDNA into the cytosol can activate innate immune pathways mediated by Toll-like receptor 9 (TLR9), Nod-like receptor NLRP3 inflammasome, and of particular relevance here, the cGAS-STING pathway(17).

The cGAS-STING pathway is an immune response that recognizes cytosolic DNA, which can come from a variety of sources including the mitochondria(18–20). Upon binding to DNA, cGAS (cyclic GMP-AMP synthase) dimerizes and becomes activated, whereupon it catalyzes the synthesis of cyclic dinucleotide cGAMP (2′3′ cyclic GMP–AMP) from ATP and GTP. These cGAMP dinucleotides then bind STING (stimulator of interferon genes), causing conformational changes that lead to STING oligomerization and subsequent translocation from the endoplasmic reticulum (ER) to the Golgi apparatus(21, 22). This activated STING recruits TANK-binding kinase 1 (TBK1)(17), which causes a series of events leading to phosphorylation of interferon regulatory factor 3 (IRF3). In turn, the phosphorylated IRF3 translocates to the nucleus where it induces expression of type I interferons, interferon-stimulated genes (ISGs), and several other inflammatory mediators, pro-apoptotic genes, and chemokines(21).

The first work linking mtDNA to the cGAS-STING pathway showed that depletion of the mitochondrial DNA packaging protein TFAM led to cytosolic release of mtDNA, resulting in cGAS-STING activation(23). More recently, deficiency in the mitochondrial CLPP (Caseinolytic mitochondrial matrix peptidase proteolytic subunit) protease subunit was also shown to result in mtDNA instability and activation of type I IFN signaling through cGAS-STING(24). Meanwhile, cytosolic mtDNA release(25) and activation of cGAS-STING have also been implicated in Parkinson disease(26). Many inflammatory diseases also show activation of cGAS-STING in damaged organs, like liver (27–29), kidney (30, 31) and pancreas(32). The involvement of TFAM and CLPP in mtDNA release into the cytosol and cGAS-STING activation raises the question of whether impairment of mtDNA maintenance caused by coding variants in other proteins might have similar consequences. One intriguing candidate is TOP1MT, a topoisomerase that helps resolve crossovers and knots that can originate during maintenance of the circular mtDNA genome. TOP1MT is the only human topoisomerase that is solely present in the mitochondria(33–35), where it plays important roles in mtDNA replication, transcription, and translation(36–39). Interestingly, a wide range of autoantibodies have been described and detected in the serum of SLE patients(40, 41), including antiLdoubleLstranded DNA (antiLdsDNA)(42, 43) and autoantibodies recognizing the nuclear topoisomerase enzyme TOP1(44–47) as well as the mitochondrial TOP1MT(48).

Here, we characterize a Variant of Unknown Significance in TOP1MT, P193L, identified in a family with several members who presented with SLE and a variety of other autoimmune disorders. Notably, activation of the cGAS-STING pathway is linked to SLE(49, 50). For example, an activating mutation in STING causes lupus-like manifestations(51). To date, the connection between cGAS-STING activation in SLE has only been noted in the context of nuclear DNA present in the cytosol(50, 52). Meanwhile, inhibition of VDAC-mediated cytosolic mtDNA release was beneficial in a mouse model of SLE(16), suggesting a role for cytosolic mtDNA in promoting SLE. Here, we uncover a novel connection between TOP1MT dysfunction and activation of the cGAS-STING pathway via mtDNA, which may contribute to the autoimmune phenotypes in several members of this family, and provide further evidence for the potential role of mtDNA in the pathogenesis of autoimmune disorders.

## Results

The family was originally ascertained on the basis of congenital leptin deficiency in the index patient’s daughter(53). However, the proband for the lupus phenotype is III-8 (the mother) has been diagnosed with hypothyroidism, systemic lupus erythematosus, Sjogren’s syndrome, fibromyalgia with peripheral neuropathy. Now in her mid 50s, she has moderate to severe pulmonary fibrosis. She is thought to have celiac disease as well, though the latter is not biopsy-proven. As seen in the extended family pedigree (Fig S1), multiple family members were diagnosed with autoimmune diseases. One of the proband sisters III-13 (from a sibship of 8) was diagnosed with lupus at age 30 years, and one of her nieces V-6 (the daughter of one of her brothers) was diagnosed with rheumatoid arthritis in her late teens. The proband husband, III-7, is clinically unaffected, though his sister III-5 was diagnosed with polyarthritis in her 40s, and one of her daughters IV’- 2 was diagnosed with lupus at age 14 years and died at age 24 years. Furthermore, another of her daughters IV’-4 was diagnosed with rheumatoid arthritis in infancy. Attempts to collect and analyse DNA on these individuals have not yet been successful. Additional clinical information on the nuclear family is available in supplemental information (Sup File 1).

### Variant Identification

Chromosomal microarray studies did not identify any microdeletions or microduplications that met clinical criteria for being considered pathogenic or likely pathogenic. Exome sequencing in members of the proband’s nuclear family did not identify any known pathogenic variants in recognized Mendelian disorders. However, TOP1MT was flagged as a potential genetic cause of monogenic systemic lupus erythematosus (SLE) when a P193L missense variant was discovered in the consanguineous family, with multiple members affected by pediatric-onset autoimmune disease, including SLE (Fig 1a). The P193L variant is a result of a point mutation at cDNA position 578C>T (NM_052963.2) that results in a proline to leucine change at amino acid 193, which is predicted to be possibly damaging and deleterious with minor allele frequency (MAF) of 0.00002786 in Genome Aggregation Database v2.1.1, and a MAF of 0.0001307 within the South Asian superpopulation sampled by v2.1.1. The identified variant occurs at a highly conserved site in the core domain of TOP1MT (Fig 1b), which is located within the DNA binding barrel (Fig 1c).

**Figure 1.**
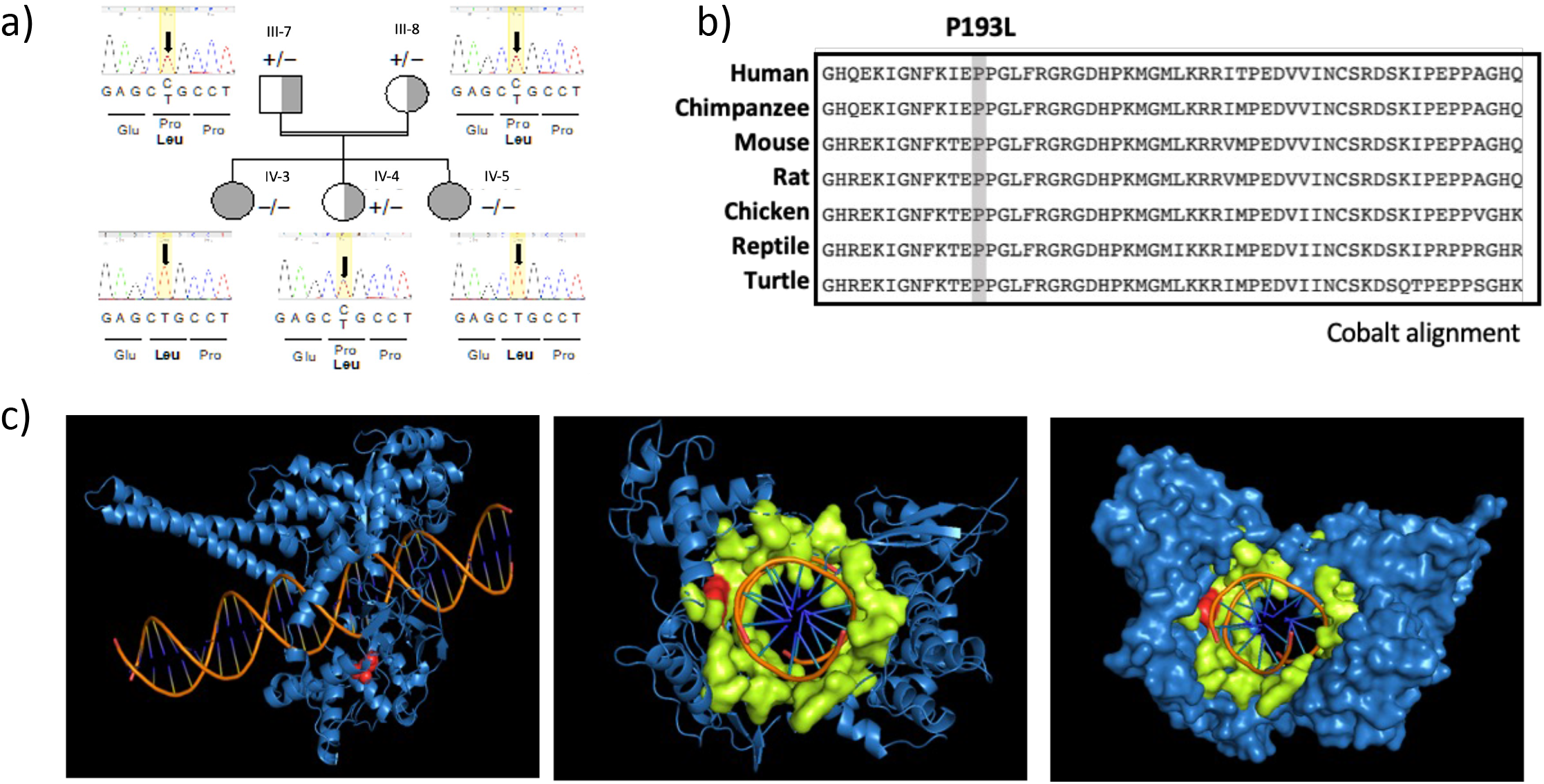
Family with the P193L TOP1MT variant and protein structure. **a)** Pedigree of the proband’s nuclear family indicating the P193L genotype; the sequences show the nucleotide identity at position C578T where a missense Pro193Leu TOP1MT alteration was found. An extended family pedigree can be found in Figure S1. **b)** Amino acid alignments of TOP1MT proteins from a variety of vertebrate species generated using COBALT shows the conservation of the amino-acid variant identified in the affected family. **c)** Structural modelling of the TOP1MT protein bound to double-stranded DNA. Residues in the nucleic acid binding barrel are shown in lime, with the modelled P193L variant shown in red.

Preliminary functional studies in patient-derived white blood cells from family members that were either heterozygous or homozygous for the P193L variant showed impaired oxidative phosphorylation (OxPhos) (Fig S2) compared to healthy controls, suggestive of underlying mitochondrial dysfunction. However, additional work was necessary to confirm whether the P193L variant impacts the functionality of TOP1MT, and to investigate potential mechanisms that might link TOP1MT to SLE.

### TOP1MT depletion leads to cytosolic mtDNA release and activation of cGAS-STING signaling

Given the SLE and other autoimmune disease phenotypes in the family, and the role of TOP1MT as a mediator of mtDNA, we hypothesized that TOP1MT dysfunction could contribute to cytosolic release of mtDNA and activation of the cGAS-STING pathway. To investigate this notion, we took advantage of HCT116 cells wherein TOP1MT is knocked out via CRISPR- Cas9(54). We first measured the abundance of mtDNA present in cytosol extracts. Using several primer sets that amplify distinct regions of the mtDNA, quantitative PCR consistently detected significantly higher levels of mtDNA in the cytosol extracts of HCT116 *TOP1MT* KO cells (HCT116 KO) compared to HCT116 wild type (HCT116 WT) (Fig 2a). Fluorescence *in situ* hybridization (FISH) also confirmed higher levels of cytosolic mtDNA in HCT116 KO cells compared to HCT116 WT (Fig 2b&c). Moreover, the cytosolic mtDNA in HCT116 KO cells colocalized with cGAS (Fig 2b&d), implicating the cGAS-STING innate immune pathway.

**Figure 2.**
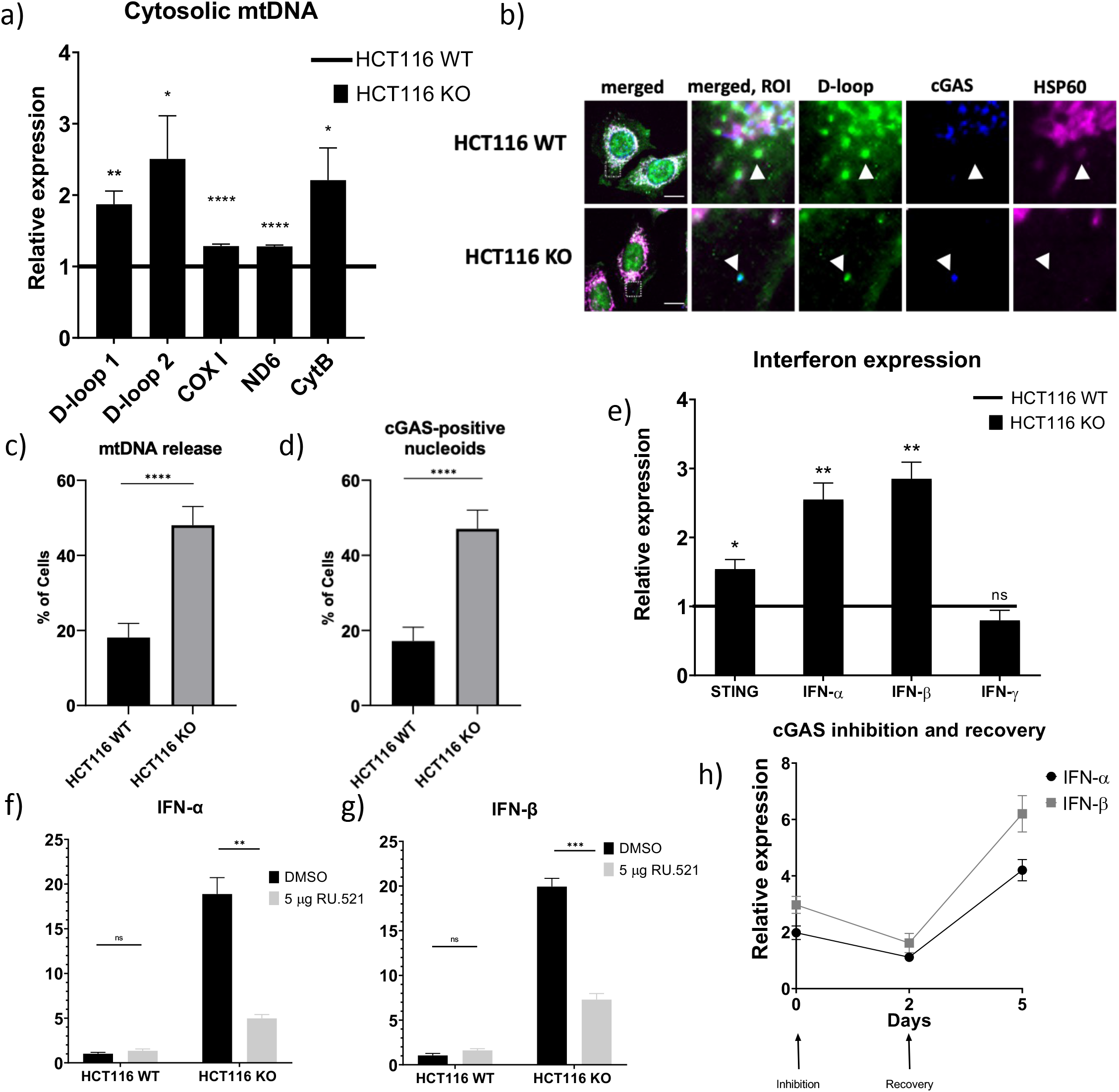
Elevated cytosolic mtDNA release and cGAS-STING pathway activation in TOP1MT KO cells. **a)** Cytosolic mtDNA was quantitated via qPCR using the indicated primer sets and was found to be elevated in TOP1MT KO cells compared to WT cells. Normalization was done relative to the corresponding amplicons in total DNA. **b)** Confocal images of mtDNA FISH (D-loop probe, green), followed by immunofluorescence against cGAS (blue) and HSP60 (magenta). Scale bar = 10 um. **c)** Quantification of the number of cells with non-mitochondrial nucleoids (N = 105 for HCT116 WT, N = 102 for HCT116 KO; pooled from 3 replicate experiments). **d)** Quantification of number of cells exhibiting colocalization of non-mitochondrial nucleoids with cGAS, as in (b). **e)** qRT-PCR analysis of STING, type I and type II interferons expression in TOP1MT KO cells compared to WT cells. Inhibition of cGAS by 5 µg/ml RU.521 for 24 hrs lowers type I interferon expressions in TOP1MT-KO cells for both IFN- ⍺ **(f)** and IFN-β**(g)**, as measured by qRT-PCR. **h)** Quantification of type I interferon levels by qRT-PCR in TOP1MT KO cells following 24 hours treatment with 5 µg/ml RU.521 and a 5 day recovery in fresh media. All statistical analysis were done using unpaired student t-test and p values * <0.05, ** <0.01, ***<0.001 and **** <0.0001. ‘ns’ signifies no significant differences between indicated groups. Error bars represent standard error of mean.

Next, to see if the increased cytosolic mtDNA activated cGAS-STING signaling, we examined the expression of type I interferons. Consistent with activation of the cGAS-STING signaling pathway, both interferon-alpha (IFN-α) and interferon-beta (IFN-β) were elevated in HCT116 KO cells. However, levels of the type II interferon-gamma (IFN-γ), which is not regulated by cGAS-STING, were not altered (Fig 2e). As there are multiple activators of innate immune responses, we wanted to confirm that this type I interferon elevation occurred through the cGAS-STING pathway. To this end, 24 hr treatment of cells with RU.521, an inhibitor of cGAS(55), led to lower interferon levels among HCT116 KO cells, whereas basal levels in HCT116 WT cells remained unchanged (Fig 2f & 2g). Subsequently, upon removal of RU.521, the interferon levels were further elevated again in HCT116 KO cells relative to HCT116 WT controls (Fig 2h). Collectively, these novel findings suggest that loss of TOP1MT results in increased levels of cytosolic mtDNA, and increased cGAS-STING dependent expression of IFN-α and IFN-β transcripts.

Although we saw a significant increase in cGAS-dependent levels of IFN-α and IFN-β in HCT116 cells, this increase was relatively mild compared to a typical type I interferon response during viral infection, likely due to the fact that HCT116 cells are a cancer cell line and have been reported to be hypo-responsive to cGAS-STING pathway activation (56, 57). Thus, we wanted to confirm our findings in additional cells lines. To this end we knocked down TOP1MT via siRNA in telomerase immortalized human fibroblasts (Fig S3a), as well as in mouse embryonic fibroblasts (MEFs) (Fig S3b). We observed a significant increase in several interferon stimulated genes (ISGs) (*e.g.*, Cxcl10, Ifit1, Ifit3) in human fibroblasts, grown at either 20% oxygen (Fig 3a) or 3% oxygen (Fig 3b), as well as in MEFs (Fig 3c). Consistent with this elevation of ISGs being dependent on cGAS-STING signaling, cGAS^-/-^ MEFs did not show the same strong elevation of ISG levels upon TOP1MT depletion (Fig 3c). Together, these findings identify a novel role for TOP1MT in mediating mtDNA release and activation of cGAS-STING innate immunity.

**Figure 3.**
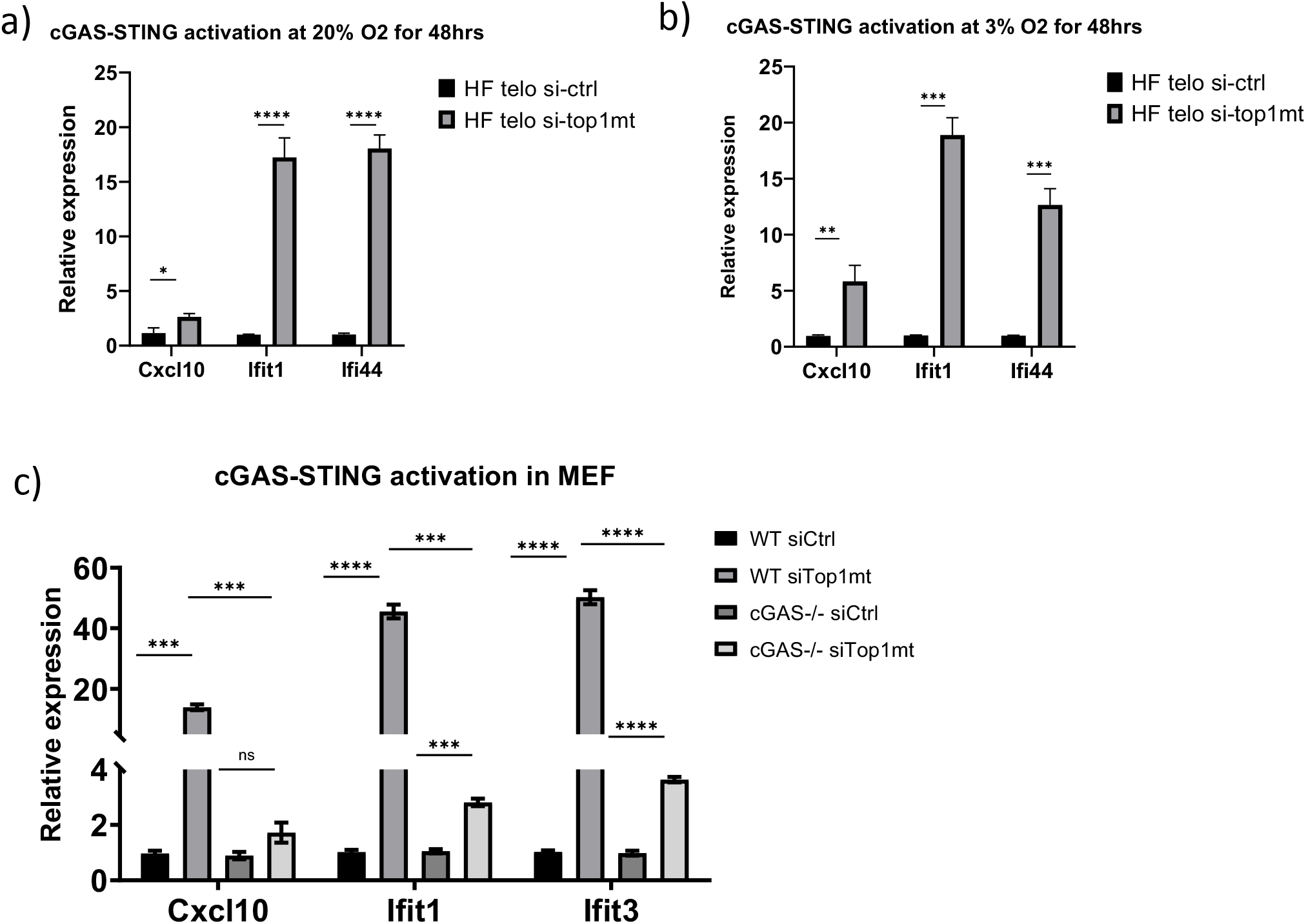
cGAS-STING pathway induction upon TOP1MT ablation in human and mouse fibroblasts. **a,b)** siRNA knockdown of TOP1MT causes strong induction of ISGs, as measured by qRT-PCR, in telomerase immortalized human fibroblasts at both 20% (a) and 3% oxygen (b). **c)** ISG expression measured by qRT-PCR is elevated in WT MEFs upon Top1mt knockdown, but this elevation is abrogated in cGAS knockout MEFs. All statistical analysis were done using unpaired student t-test and p values * <0.05, ** <0.01, ***<0.001 and **** <0.0001. ‘ns’ signifies no significant differences between indicated groups. Error bars represent standard error of mean.

### Generation of TOP1MT Rescue Cell Lines

Given the mitochondrial dysfunction noted in patient-derived blood cells (Fig S2), we examined the ability of the TOP1MT P193L variant to rescue multiple mitochondrial parameters impaired in HCT116 KO cells(39). As described previously(39), starting from HCT116 WT and HCT116 KO, we generated four different stable cell lines using retroviral constructs expressing TOP1MT and the mCherry fluorescent protein separated by a self-cleavage peptide (TOP1MT- T2A-mCherry), or an empty vector control. This construct leads to expression of TOP1MT, which will be directed to the mitochondria by the mitochondrial targeting sequence, and mCherry, which can be used to monitor transduction. Two control cell lines, henceforth referred to as WT-ctrl and KO-ctrl, were generated by transducing an empty vector expressing only mCherry in the HCT116 WT and HCT116 KO cells respectively. A third cell line was generated by re-expressing wild type TOP1MT (indicated as Rescue). Finally, a fourth cell line expressing the patient variant P193L was also established (referred to as P193L).

Expression of mCherry was used to sort the positively transduced cells by flow cytometry. Transduction was further confirmed by PCR amplification of the inserted TOP1MT open reading frame (Fig 4a), and DNA sequencing to confirm the identity of the Rescue and the P193L variant cell lines (Fig 4b). Next, immunoblotting was used to confirm the expression of TOP1MT proteins (Fig 4c). A band of the predicted size of ∼70 kDa corresponding to TOP1MT protein was observed WT-ctrl, Rescue and P193L cells, but not in the KO-ctrl cells (Fig 4d). Notably, the levels of TOP1MT protein in the Rescue and P193L lines were ∼10-fold higher than the endogenous TOP1MT protein levels in the WT-ctrl cells. Further validation of the protein expression was confirmed via immunofluorescence imaging, wherein overexpressed TOP1MT protein was detected, and confirmed to localize to mitochondria (Fig 4e).

**Figure 4.**
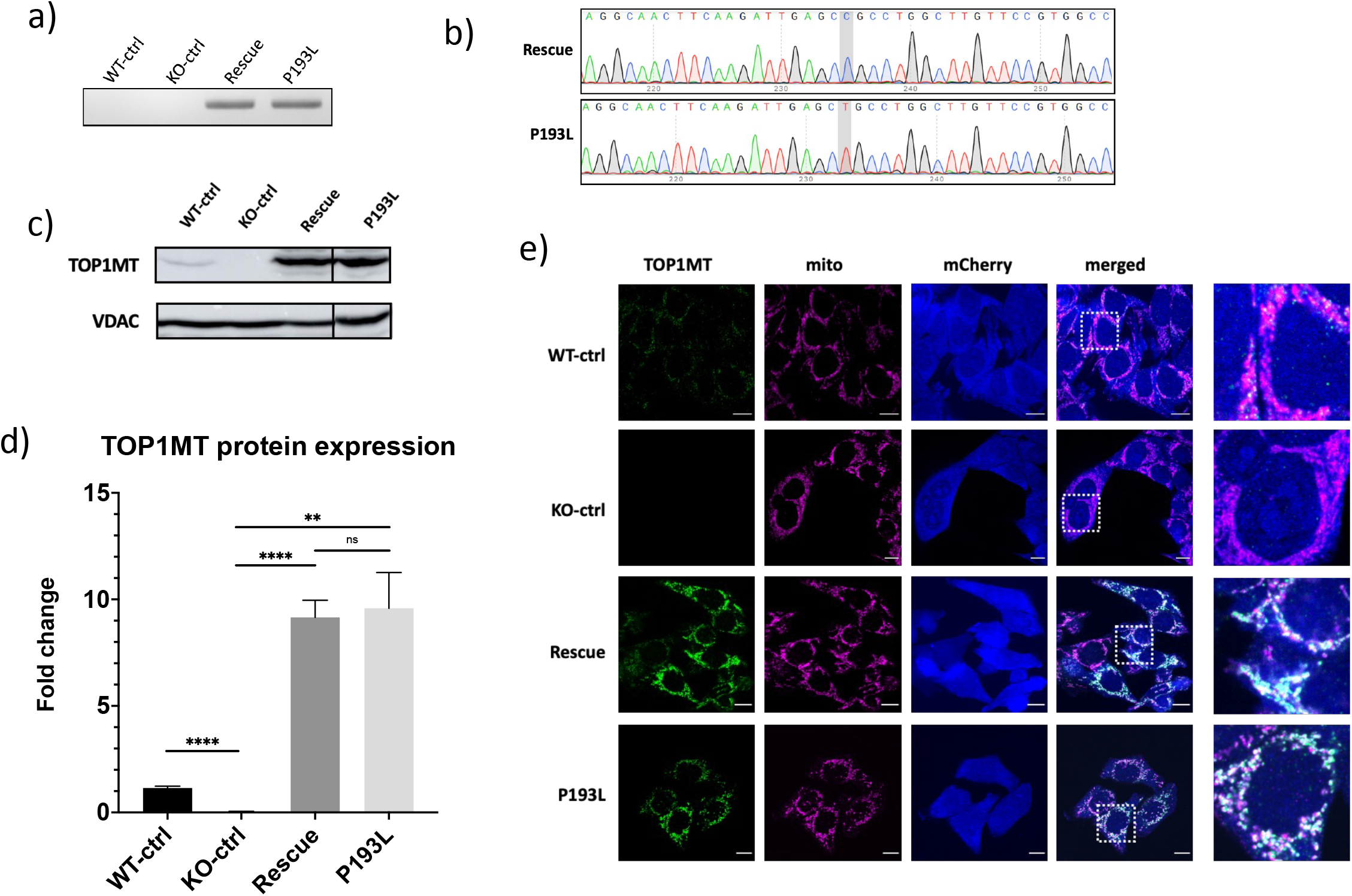
Generation of stable line expressing the variant form of TOP1MT. a) PCR amplification targeting the coding region of the viral-encoded TOP1MT open reading frame, spanning patient variant, confirms amplicons are present only in HCT116 KO cell lines stably re-expressing TOP1MT, namely the Rescue and P193L lines. **b)** Chromatograms from Sanger sequencing of the amplicons produced from (a). The 578C residues (shaded), confirm the identity of the cell lines. **c)** Western blots probed for TOP1MT antibody confirm the TOP1MT expression in stably transduced cell lines. VDAC was probed as a load control. **d)** Quantification of western blots from 4 replicates shows the fold change in protein abundance compared to VDAC control from four different replicates. All statistical analysis were done using unpaired student t-test; where error bars represent standard error of mean and p values ** <0.01, **** <0.0001 and ‘ns’ signifies no significant differences between indicated groups. Error bars represent standard error of mean. **e)** Representative confocal images of TOP1MT localization in stable lines where fixed cells were stained for immunofluorescence with anti-TOMM20 (mitochondria, magenta) and anti-TOP1MT (green) primary antibodies. Viral transduction was confirmed by the expression of mCherry (blue). The final column shows the zoom-in of the region indicated by the white dashed box in the merged column. Images confirm the mitochondrial localization of TOP1MT-P193L patient variant. Scale bar represents 10 μm. Uncropped images of the western blot are available in Fig S8b.

Next, we wanted to determine the ability of the P193L variant to rescue the aberrant functions previously identified in HCT116 KO cells(39), including: reduced nucleoid size, the hyperfused mitochondrial network, and decreases in mitochondrial transcription, mtDNA replication, mitochondrial translation and oxidative phosphorylation.

### Nucleoids

Based on our previous work(39), and the role of TOP1MT in preventing negative supercoiling of mtDNA(58) we examine mtDNA. To look at supercoiling, we used a southwestern blot approach that labels mtDNA with BrdU, which can then be separated on an agarose gel, transferred to a membrane and visualized by immunoblotting. To confirm which bands represent linear, relaxed, or supercoiled mtDNA, DNA was treated with either BamHI or topoisomerase enzymes. Consistent with the expected reduction of topoisomerase activity, we found that KO-ctrl cells had a higher ratio of supercoiled to relaxed mtDNA compared to WT-ctrl cells, and that this was reversed in the Rescue cells (Fig 5a). Though the P193L line appeared to show increased supercoiling of mtDNA relative to the rescue line, the ratio of supercoiled mtDNA to the relaxed mtDNA was not significantly altered (Fig 5b).

**Figure 5.**
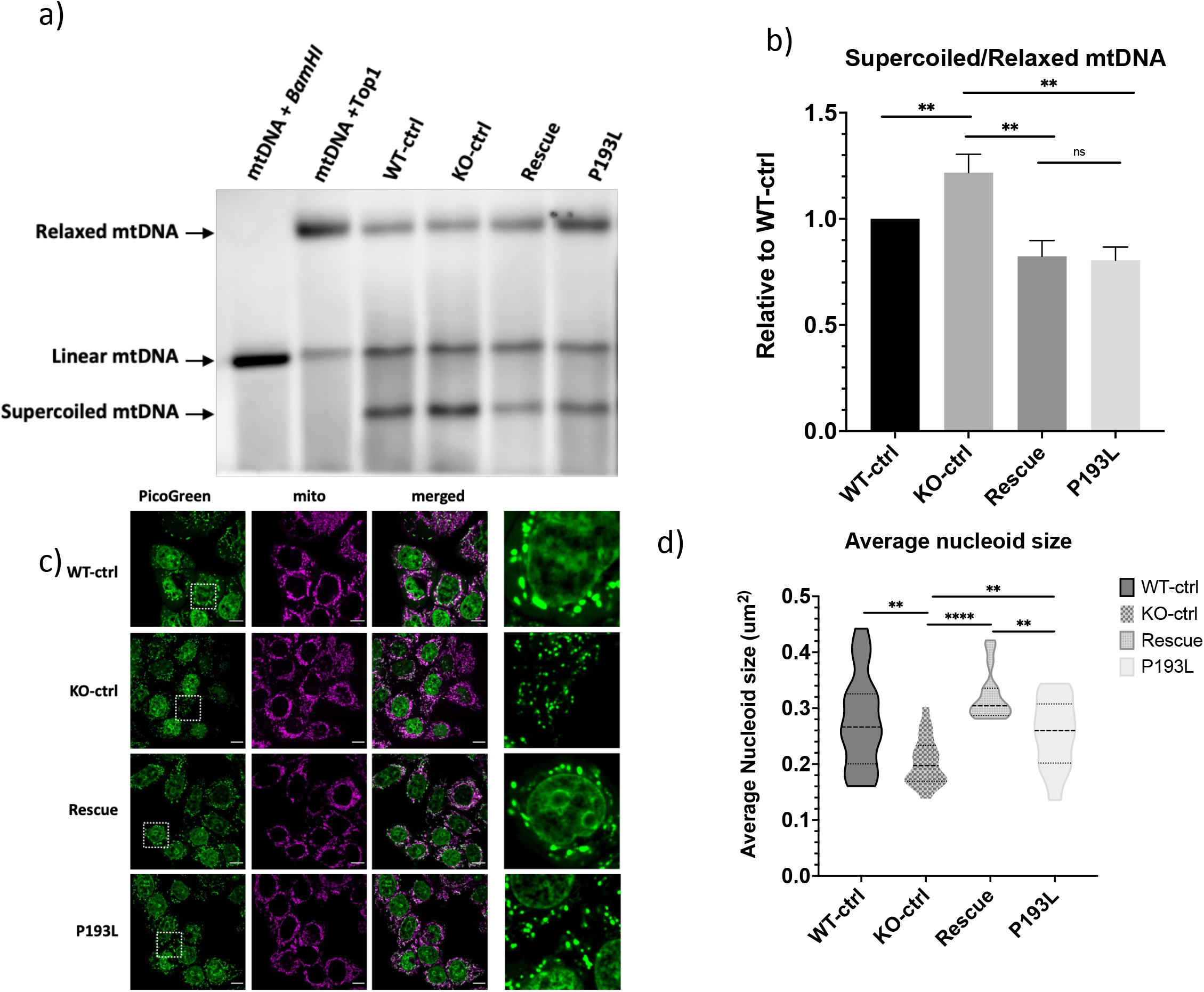
TOP1MT-KO cells stably expressing P193L alter mitochondrial genome supercoiling. a) Representative southwestern blot showing increased supercoiled mtDNA in KO ctrl cells and P193L cells. **b)** Quantification of the ratio of supercoiled to relaxed mtDNA from southwestern blots as in (a) from four independent experiments, normalized relative to control cells. **c)** Representative confocal images of live cells stained with MitoTracker Far Red (mitochondria) and PicoGreen (dsDNA: nuclear and mtDNA), where the final column shows a zoom-in of the region indicated in the white dashed box for the PicoGreen signal. Scalebars represent 10μm. **d)** Violin plots showing quantification of the average mtDNA nucleoid sizes from 25 cells stained as in (c). All statistical analysis were done using unpaired student t-test and p values ** <0.01, **** <0.0001 and ‘ns’ signifies no significant differences between indicated groups. Error bars represent standard error of mean.

As an alternative approach to gain insight on the mtDNA topology, we quantified the size of individual nucleoids stained with PicoGreen using confocal microscopy (Fig 5c). As expected, and consistent with the southwestern blot analysis showing increased supercoiling, mtDNA nucleoids were significantly smaller in KO-ctrl cells, and this was reversed by overexpression of the WT TOP1MT in the Rescue line. However, the overexpression of the P193L variant failed to fully rescue the nucleoid sizes (Fig 5d) to the same level as the WT protein, suggesting that this variant has reduced topoisomerase activity.

### Mitochondrial Morphology

The mitochondrial network is dynamic and can change in response to various physiological cues. In KO-ctrl cells, mitochondrial networks imaged by confocal microscopy were more fused than their WT-ctrl counterparts, and this was reversed by re-expression of the WT protein in the Rescue cells (Fig S4a & b). However, expression of the patient variant had an intermediate effect on mitochondrial morphology, consistent with the variant being unable to completely rescue the effects of TOP1MT loss.

### Mitochondrial Transcription

We next examined the expression of mtDNA-encoded transcripts, as the supercoiling extent of mtDNA directly influences the transcription of the mitochondrial genome(37–39). While the WT TOP1MT could restore mitochondrial transcript levels, the P193L variant was slightly less efficient at doing so (Fig 6a). This reduced rescue is likely because TOP1MT alters mtDNA topology, which impacts mitochondrial transcription *in vitro*(59–63).

**Figure 6.**
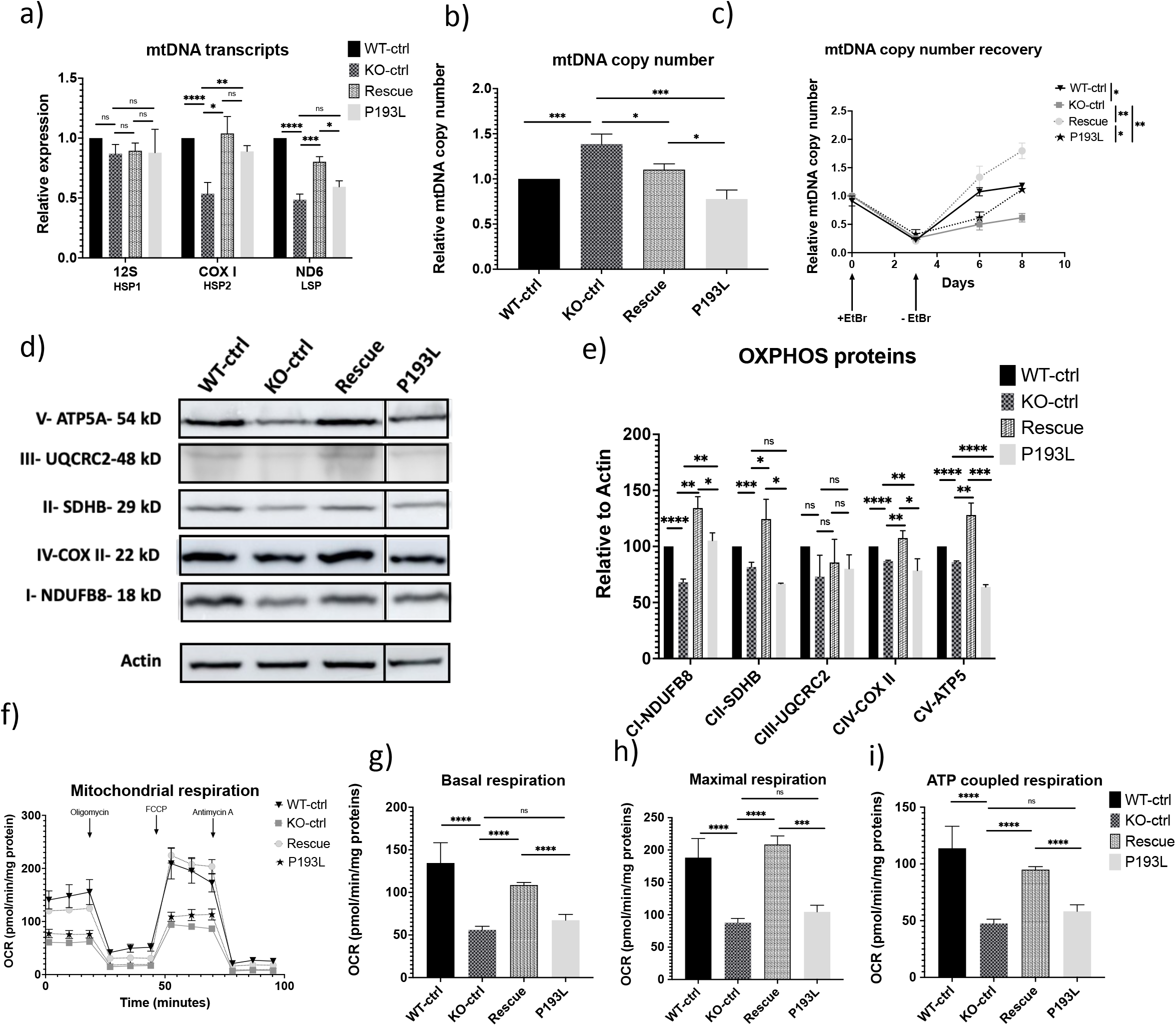
TOP1MT-KO cells stably expressing P193L affect mitochondrial DNA maintenance. a) Quantification of mtDNA transcripts from three independent biological replicates in the indicated stable cell lines via qRT-PCR for the indicated genes from the three mtDNA promoters (HSP1 (12S), HSP2 (COX I) and LSP (ND6)). **b)** Relative mtDNA copy number from three independent biological replicates, determined by qPCR relative to 18S, is rescued in P193L cells. **c)** Time course of mtDNA copy number changes (via qPCR) of ethidium bromide (EtBr) depletion (3 days, 1 µg/ml) followed by repletion (5 days, no EtBr), shows slower recovery in P193L cells compared to Rescue cells. Experiment was performed with three biological replicates. **d)** Representative western blots show reduced levels of the indicated OXPHOS complex proteins in P193L cells. Blots are cropped at the indicated sizes, with full uncropped blots available in Fig S8c. **e)** Quantification of western blots for OXPHOS proteins as in (d), from three independent experiments, corrected to Actin as a load control. Reduction in OXPHOS proteins encoded by both nuclear (i.e., NDUFB8, SDHB, UQCRC2, ATP5) and mtDNA (i.e., COXII) genomes was observed in P193L cells lines. **f)** Oxygen consumption rates (OCR) analyzed in the indicated cell lines using the Seahorse XF24 extracellular flux analyzer. The OCR data was used to calculate basal respiration **(g)**, maximal respiration **(h)** and ATP production **(i)**. Data presented is from a representative experiment with 5 technical replicates. All statistical analysis were done using unpaired student t-test and p values * <0.05, ** <0.01, *** <0.001 and **** <0.0001. ‘ns’ signifies no significant differences between indicated groups. Error bars represent standard error of mean.

### mtDNA copy number and replication

In addition to the effect on transcription, we wanted to determine the ability of the P193L variant to reverse the elevated mtDNA copy number (Fig 6b) and reduced mtDNA replication rates (Fig 6c) observed in KO-ctrl cells. In this regard, the P193L variant was able to rescue steady-state mtDNA copy number levels to a similar degree as the WT protein. To look at mtDNA replication rates, we monitored the rescue of mtDNA copy number following depletion by ethidium bromide. While the P193L variant showed improved mtDNA repletion compared to the HCT116 KO cells, with nearly complete rescue by day 8, mtDNA replication was significantly slower than that in cells re-expressing the WT TOP1MT protein, again indicative of reduced functionality.

### Expression of Mitochondrial Proteins

As TOP1MT can also impact the translation of mitochondrial proteins independently of its topoisomerase activity and changes in transcripts(39), we studied the effect of P193L on the expression of oxidative phosphorylation (OxPhos) proteins encoded by both the nucleus (NDUFB8, SDHB, UQCRC2 and ATP5) and mtDNA (COXII) (Fig 6d). Notably, global OxPhos protein levels can reflect mitochondrial translation, as the levels of nuclear-encoded OxPhos proteins depend on the expression of mtDNA encoded subunits(64). Expression of Complex I subunits NDUFB8, Complex II subunit SDHB, Complex III subunit UQCRC2, Complex IV subunit COXII and Complex V subunit ATP5A (Fig 6e) showed similar trends with reduced expression in P193L cells compared to Rescue. Although we did not see a significant decrease in SDHB and UQCRC2 in these experiments, the results were trending in the same direction. Combined, these results indicate that the P193L variant is unable to completely restore reduced OxPhos proteins expression.

### Mitochondrial Respiration

Following up on the changes in OxPhos proteins, we next investigated mitochondrial respiration in our stable cell lines (Fig 6f) to examine if the reduced expression of OxPhos proteins affect the oxygen consumption and mitochondrial energy production. As seen previously(39, 65), we found that basal respiration (Fig 6g), maximal respiration (Fig 6h), and ATP-coupled respiration (Fig 6i), were all reduced in the KO-ctrl cells, and restored by re-expressing the WT TOP1MT protein. However, the cells re-expressing the P193L patient variant showed no rescue at all relative to the KO-ctrl cells. Notably, this finding is consistent with patient-derived white blood cells showing impaired OxPhos (Fig S2).

### cGAS-STING pathway in rescue cell lines

Finally, we examined the effect of the P193L variant on the novel cGAS-STING pathway activation that we described above in the HCT116 KO cells. For this purpose, we started by looking at the cytosolic mtDNA levels. Unexpectedly, both lines stably overexpressing either WT or P193L forms of TOP1MT rescued the cytosolic mtDNA levels to the same extent (Fig S5a-c). Similarly, qRT-PCR analysis of type I interferon expression was rescued in HCT116 KO cells overexpressing either WT or P193L forms of TOP1MT (Fig S5d). Meanwhile, high level expression of other TOP1MT variants linked to cardiomyopathy (Fig S5a-b), namely R198C and V338L, also rescued the cytosolic mtDNA (Fig S5c) and the interferon levels (Fig S5d). However, these lines express TOP1MT protein at approximately 10-fold above endogenous levels, which we reasoned could be compensating for a loss of activity in the P193L protein.

### Characterization of low re-expression of P193L and WT TOP1MT variants

To avoid the complications of high level TOP1MT protein expression, we used flow cytometry to sort for cells expressing relatively low levels of mCherry, which should correlate directly to TOP1MT protein expression. As shown by western blot (Fig 7a), we were able to select a population of cells expressing TOP1MT protein much closer to endogenous levels (Fig 7b). These lines were labeled ‘Rescue low’, and ‘P193L low’. We first looked at nucleoid size as a proxy for mtDNA supercoiling, as this assay is more sensitive than the southwestern assay(39). We found that low expression of the P193L variant was unable to rescue nucleoid size to any degree (Fig 7c-d), compared to the partial rescue when the P193L variant was expressed at a high level (Fig 5d).

**Figure 7.**
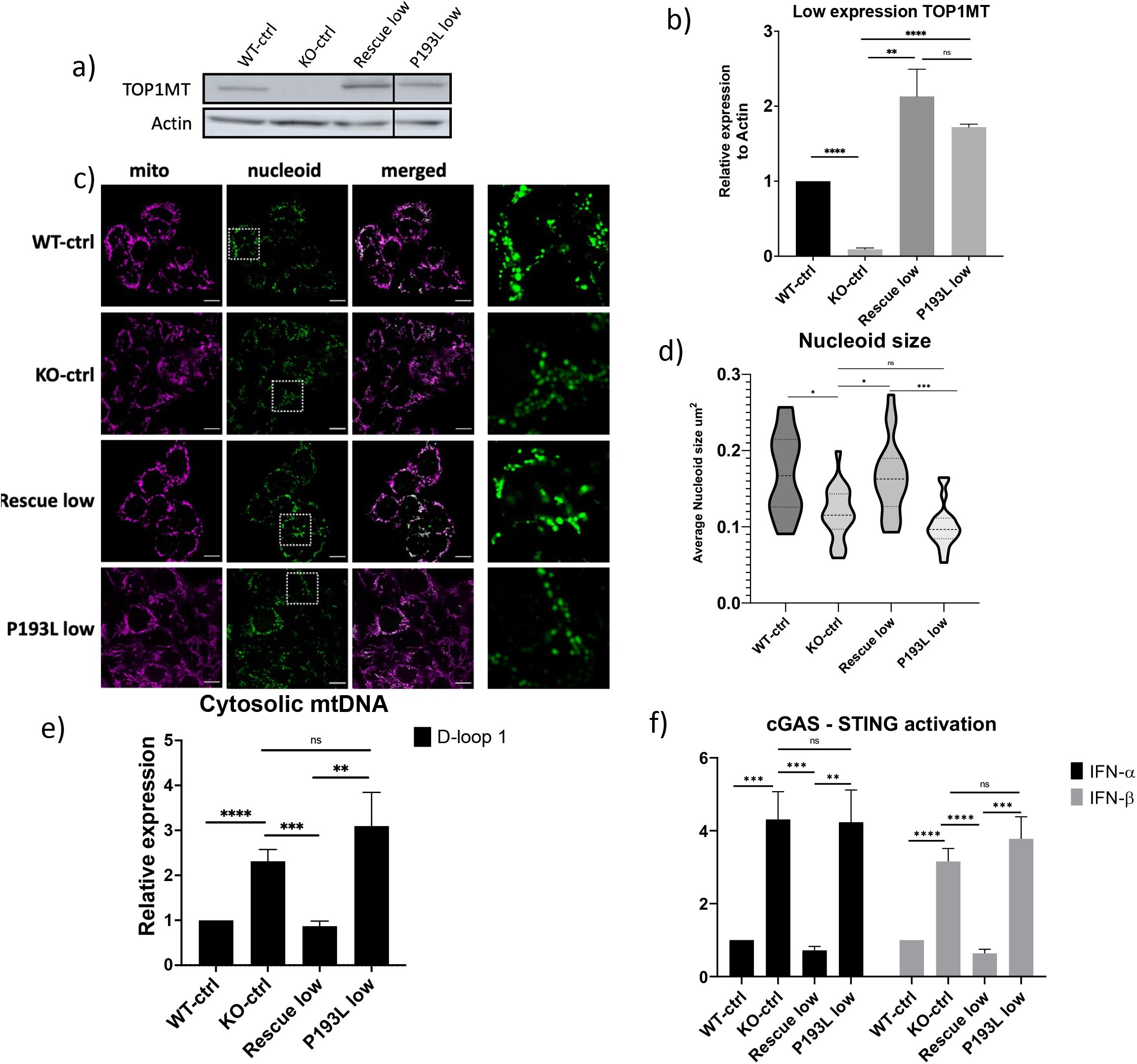
TOP1MT-KO cells stably expressing low levels of P193L do not rescue cytosolic mtDNA levels, reduced nucleoid sizes, or elevated interferon transcripts. a) Representative western blots show near endogenous expression of the TOP1MT protein in cells sorted for lower expression of both WT and P193L variants. b) Quantification of western blots of TOP1MT protein as in (c), from three independent experiments, corrected to Actin as a load control. c) Representative confocal images of live cells stained with MitoTracker Far Red (mitochondria) and PicoGreen (dsDNA: nuclear and mtDNA) where the final column shows the zoom-in of the region indicated in the white dashed box for the PicoGreen signal. Scale bars represent 10μm. d) Violin plots showing quantification of the average mtDNA nucleoid size from 25 cells stained as in (c). e) Quantification of cytosolic mtDNA levels from cells as in (a) via qPCR using the indicated primer set. Normalization was done relative to the corresponding amplicons in total DNA from the same cells. f) qRT-PCR analysis of type I interferon expression from cells as in (a) show elevated type I interferon levels in cells expressing endogenous levels of the P193L variant. All statistical analysis were done using unpaired student t-test and p values * <0.05, ** <0.01, *** <0.001 and **** <0.0001. ‘ns’ signifies no significant differences between indicated groups. Error bars represent standard error of mean. Uncropped images of western blots are available in Fig S8d.

Next, we analyzed the cytosolic mtDNA levels from these cells. While the low-level expression of TOP1MT WT was still able to rescue the cytosolic mtDNA levels, the low-level expression of the P193L variant was unable to provide rescue (Fig 7e). In a similar fashion, the type I interferon levels remained elevated in cells expressing low levels of the P193L variant but were restored to normal in cells expressing low levels of the WT protein (Fig 7f). However, low-level expression of R198C and V338L TOP1MT variants linked to cardiomyopathy(39) (Fig S6a-b), fully rescued the nucleoid size (Fig S6c), the cytosolic mtDNA (Fig S6d), and the interferon levels (Fig S6e), suggesting that the cGAS-STING activation is specific to the P193L variant and pathogenic autoimmune activation. Notably, low expression of P193L also failed to show any rescue of mitochondrial morphology, while low expression of R198C and V338L showed partial rescue (Fig S4c &d).

As another means to examine activation of the cGAS-STING pathway, we used immunofluorescence to monitor the cellular localization of STING, as previous reports show that activated STING translocates from the ER to the Golgi(66). Notably, there is a stronger co-localization signal between STING and the ER in both WT-ctrl and Rescue low lines, compared to the KO-ctrl and P193L low lines (Fig 8a&b). Conversely, there were higher levels of colocalization between STING and Golgi in the KO-ctrl and P193L low cells, compared to WT-ctrl and Rescue low cell line (Fig 8c&d). These findings are consistent with the P193L variant being unable to rescue cGAS-STING signaling in absence of TOP1MT. Meanwhile, both R198C low and V338L low variants showed a localization pattern similar to the rescue line (Fig S8a and b). Altogether, our data provide support for the notion that the P193L variant has impaired functionality, which leads to activation of the cGAS-STING innate immune pathway via promoting the cytosolic release of mtDNA.

**Figure 8.**
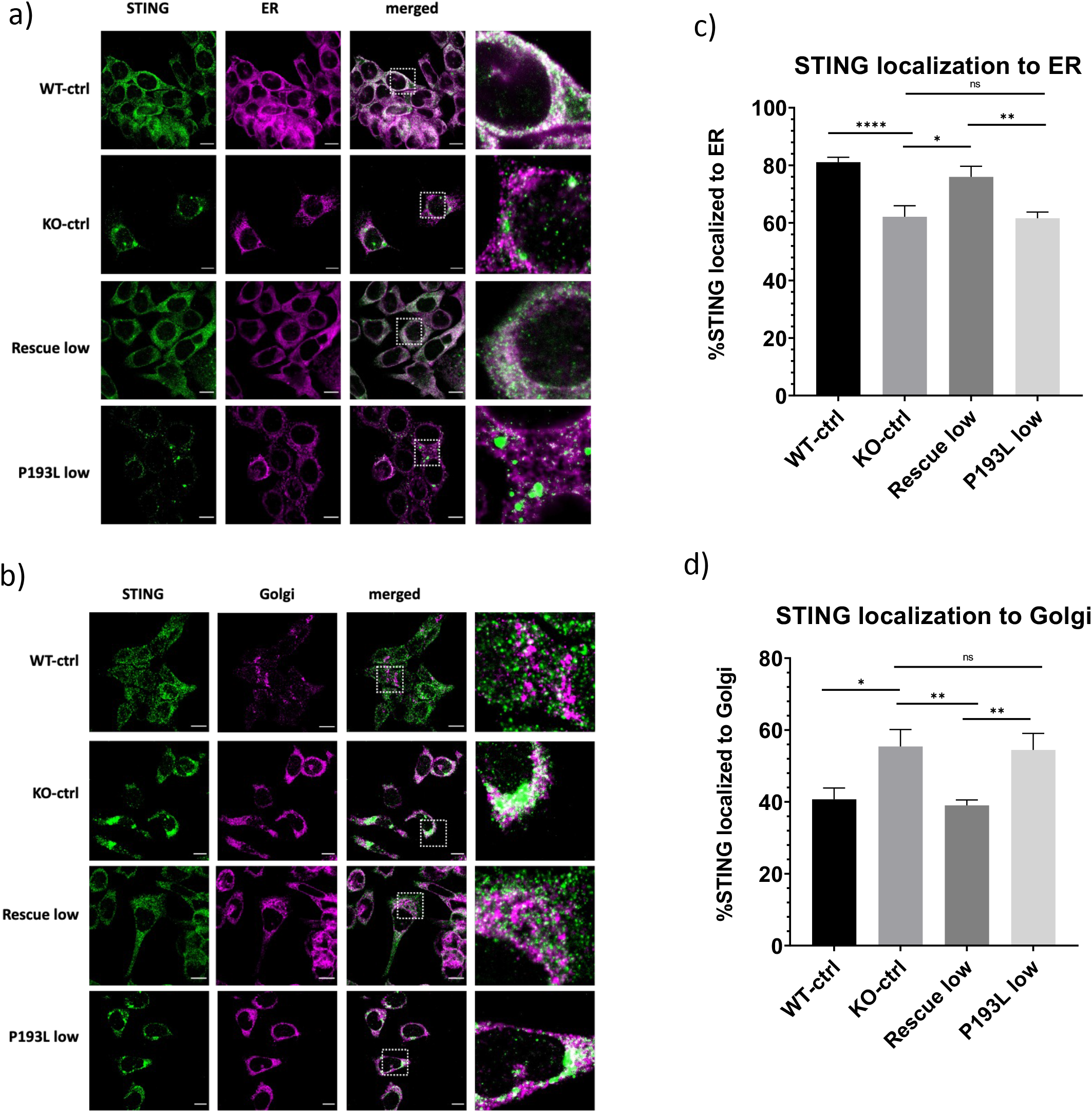
STING relocates from the ER to the Golgi in TOP1MT KO and P193L low cells. Representative confocal images of fixed cells to visualize the co-localization (white) of STING (green) with either the endoplasmic reticulum (ER) or Golgi (magenta). a) Cells stained with primary antibodies for immunofluorescence with anti-STING (green) and anti-Calnexin (ER, magenta. b) Cells stained with primary antibodies for immunofluorescence with anti-STING (green) or anti-TGN46 (Golgi, magenta). The final column shows the zoom-in of the region indicated by the white dashed box in the merged column. Scale bar represents 10 μm. Localization of STING to the Golgi, and the absence of STING in the ER, which are indicative of STING activation, were observed in the TOP1MT KO and P193L low lines and confirmed by quantification of the percentage of STING signal co-localizing to the ER (c) or Golgi (d) from at least 10 images as in (a) and (b) respectively. All statistical analysis were done using unpaired student t-test and p values*<0.05, ** <0.01 and **** <0.0001. ‘ns’ signifies no significant differences between indicated groups. Error bars represent standard error of mean.

## Discussion

The discovery of the novel TOP1MT missense P193L variant present among multiple family members with autoimmune phenotypes led us to investigate whether TOP1MT could impact mtDNA-mediated activation of the cGAS-STING innate immune response. We found that the loss of TOP1MT in HCT116 cells led to increased levels of cytosolic mtDNA and activation of the cGAS-STING pathway, as reflected by increased levels of type I interferons. Demonstrating that this pathway is conserved in multiple cell types and organisms, we found that depletion of TOP1MT in both human fibroblasts and mouse embryonic fibroblasts also resulted in activation of cGAS-STING signaling. To show that the interferon signaling is dependent on the cGAS-STING pathway, we found that the cGAS inhibitor RU.521 blunts signaling in HCT116 TOP1MT KO cells, while genetic ablation of cGAS also prevented the signaling in MEFs depleted of TOP1MT.

Having established a novel link between TOP1MT loss of function and mtDNA-mediated cGAS-STING activation, we also investigated the impact of three TOP1MT variants on this pathway. We found that when expressed close to endogenous levels, the P193L variant was unique among the three TOP1MT variants we interrogated in its inability to rescue the mtDNA release and cGAS-STING activation phenotype. This finding supports the idea that the P193L variant may contribute to the autoimmune phenotypes in our family of interest, as the other variants, which are not linked to autoimmune phenotypes, were able to rescue the cGAS-STING activation. This finding also highlights the fact that rare variant burden testing for correlation of genotypes to phenotypes in TOP1MT must be supported by functional studies, as variants that do not exhibit the relevant functional properties will dilute the signal that is conferred by variants that do impact disease relevant functions.

Our findings show that the P193L variant is not fully functional. In addition to the inability of low levels of the P193L variant to reverse cGAS-STING activation in cells lacking TOP1MT, we found that it was unable to rescue several other mitochondrial phenotypes linked to TOP1MT function, even when expressed at high levels (summarized in Table 3). Importantly, the fact that the P193L variant was overexpressed by ∼10 fold in our cellular model system, likely means that we are overestimating the partial functionality of this variant. Nonetheless, high expression of the P193L variant was only capable of partially rescuing nucleoid size, transcript levels, and mtDNA copy number recovery. Similarly, mitochondrial morphology was also only partially rescued by expression of the P193L variant. Although the mechanistic connection between TOP1MT and mitochondrial morphology remains unknown, it is tempting to speculate that this likely also involves the role of TOP1MT in regulating mtDNA. Meanwhile, the P193L variant was unable to restore steady state levels of OxPhos proteins, as well as several parameters of mitochondrial respiration. This finding suggests that the role of TOP1MT in mitochondrial translation, which is distinct from its topoisomerase activity, likely contributes to the reported mitochondrial dysfunction in patient blood cells. The finding that translation is more severely impaired than topoisomerase activity for the P193L variant is similar to previous work for two other TOP1MT variants, R198C and V338L(39).

Intriguingly, the correlation between lack of topoisomerase activity, mtDNA release and elevated cGAS-STING signaling suggests that the topoisomerase activity of TOP1MT is required to prevent mtDNA release, rather than the other characterized functions of TOP1MT in transcription or translation. In this context, it is notable that R198C and V338L TOP1MT variants have impaired transcription or translation activities, but retain greater topoisomerase activity, and are able to rescue mtDNA release and cGAS-STING activation. These observations further support the notion that the transcription and translation functions of TOP1MT are not involved in mtDNA release. Nonetheless, we cannot exclude the possibility that some other, perhaps unrecognized, function of TOP1MT plays a role in preventing mtDNA release into the cytosol. Moreover, exactly how the P193L variant mechanistically leads to cytosolic release of mtDNA remains unknown. Another unresolved question is how does the P193L variant impact protein activity. In this regard, it may be notable that the P193L variant is located within the core barrel of TOP1MT, which is predicted to bind nucleic acids.

Although *TOP1MT* is not yet formally annotated to the clinome as a disease gene, increasing evidence suggests that some functional *TOP1MT* variants may indeed be pathogenic(39, 67). For example, an *in vitro* study showed that some TOP1MT variants have reduced DNA relaxation activities, including E168G, which has been described in a patient with mitochondrial deficiency syndrome(68). Meanwhile, two other TOP1MT variants were identified in a patient with a pediatric cardiomyopathy(39). The fact that the P193L variant we describe here was identified within an extended pedigree that had multiple members affected with SLE and other autoimmune diseases suggests that this is a novel pathogenic phenotype linked to TOP1MT dysfunction. However, it is not possible to rule out polygenic contributions from other genes, as no comprehensively diagnostic “lupus panel” exists to this end. To date, there are only a limited number of reports of potential pathogenic TOP1MT variants, which are associated with diverse pathogenic phenotypes(39, 67–69). As such, more examples with shared phenotypes need to be identified before TOP1MT can officially be definitively considered as a disease gene. Nonetheless, future studies looking to establish the genetic cause of SLE should also consider variants in TOP1MT. Similarly, variants in other genes encoding proteins linked to mtDNA release (*e.g.*, TFAM, POLG, CLPP, ATAD3B) could also be considered in autoimmune diseases (23, 24, 70, 71).

Genetic, epigenetic or environmental risk factors have been implicated as underlying causes of SLE(72); however, the underlying mechanism behind the disease is not always clear. With respect to whether the P193L variant contributes to the SLE phenotype in the proband and her family members affected with lupus and other autoimmune diseases, it is important to consider the already-established links between SLE and mitochondrial dysfunction. For example, general mitochondrial dysfunction and aberrant oxygen metabolism have a role in many autoimmune diseases, including SLE(73–75). Meanwhile, links between SLE and the cGAS-STING innate immune response are also important in the context of considering the contribution of TOP1MT dysfunction to SLE. Consistent with the notion the cGAS-STING activation contributes to SLE, previous reports have noted cGAS-STING activation in a subset of SLE cases(76, 77). Some of this previous work implicated nuclear DNA as the cytosolic dsDNA trigger, as studies identified impaired function of the nuclear DNA repair machinery can lead to SLE(78–84). However, it is worth noting that much of the mitochondrial DNA repair machinery is shared with the nuclear repair machinery. Thus, it is likely that the impairments in nuclear DNA repair enzymes that are linked to SLE also impact mtDNA. The fact that some SLE patients exhibited increased levels of mtDNA damage(85) supports this notion and argues that mtDNA damage may contribute to SLE. Moreover, it is notable that topoisomerases, including TOP1MT, can participate in DNA repair(86, 87). Collectively, this previous work linking SLE, mitochondrial dysfunction, and the cGAS-STING pathway, support the argument that the P193L variant in TOP1MT could contribute to SLE.

Another intriguing connection between SLE and TOP1MT is that fact that autoantibodies recognizing the nuclear topoisomerase enzyme TOP1 have been described in patients with SLE(44–47). Notably, these TOP1 autoantibodies recognize the highly conserved catalytic domain of TOP1(88), which is conserved with TOP1MT. Given the novel connection that we describe here between TOP1MT and SLE, it is tempting to speculate that these autoantibodies are in fact made against TOP1MT and cross-react with TOP1, even though they were first shown to bind TOP1. This notion is supported by the fact that SLE sera recognize TOP1MT(89). Given that 19% of autoantibodies collected from patients with SLE were able to detect TOP1MT(48), it is possible that TOP1MT plays a much wider role in SLE than currently appreciated. For example, if TOP1MT released from mitochondria along with mtDNA, these autoantibodies could also contribute to an elevated immune response contributing to SLE. Thus, beyond our current study, there is considerable evidence supporting the argument that TOP1MT plays a role in SLE.

The novel connection we describe here between TOP1MT, cGAS-STING and SLE has important clinical implications, as identifying novel SLE candidate genes such as TOP1MT, and understanding the mechanistic underpinnings of mtDNA-mediated inflammation, will be important for personalized medicine approaches. As we learn more about the different causes of SLE and their underlying mechanisms, this will provide a clearer picture of which treatments might be beneficial. For example, cGAS inhibitors like RU.521 may prove to be effective in patients where the cGAS-STING pathway is elevated. Meanwhile, it is also important to consider the effects of drugs that impact mtDNA, a key example being the chemotherapeutic agent doxorubicin, which causes mtDNA damage, cytosolic release of mtDNA, and activation of the cGAS-STING pathway(68, 90, 91). While cGAS-STING activation by doxorubicin may be advantageous by activating innate immune signaling in cancer(92), it could be detrimental in other circumstances by activating the immune response inappropriately. In this regard, loss of TOP1MT increases sensitivity to doxorubicin treatment [76], suggesting TOP1MT variants could be relevant to the inflammatory cardiotoxicity associated with this commonly used cancer drug.

### Conclusions

In summary, this report provides the first evidence linking TOP1MT to release of mtDNA to the cytosol, activation of the innate immune response through the cGAS-STING pathway, and ultimately increased levels of interferons. Our findings also support the notion that the P193L variant could contribute to the autoimmune phenotypes in the patients, and provide novel mechanistic insight how mitochondrial dysfunction might contribute to autoimmune diseases via mtDNA-mediated activation of innate immunity. A better understanding of the causes of autoimmune diseases will help to elucidate the complex heterogeneity of these diseases, and could lead to novel therapeutic approaches.

## Materials and Methods

### Mutation identification

In accordance with the principles of the Declaration of Helsinki, all study participants provided informed consent to next-generation sequencing as part of the BCCHRI study “Comprehensive Characterization of Genetic Disorders of Body Weight” (UBC CREB Study approval number H08-00784).

### Cell culture

Primary mouse embryonic fibroblasts (MEFs) were generated from E12.5–14.5 embryos. Cells were maintained in DMEM (D5756, Sigma-Aldrich) with 10% FBS (97068-085, VWR). McCoy’s 5A (modified) media (Gibco, 16600-82) containing l-Glutamine and supplemented with 10% fetal bovine serum (FBS) was used to culture HCT116 control (HCT116 WT), HCT116 TOP1MT-KO (HCT116 KO) and stably transfected cells. Cells were seeded at 1.5 x 10^6^ cells seeding density in 10cm plates and allowed to grow for two days before being harvested for analysis. DMEM (Gibco, 11965092) supplemented with 10% FBS was used for Phoenix cells for the retrovirus generation, MEFs and HFF-1 cells. All cells were maintained at 37 °C and 5% CO_2_.

### Plasmids & cloning

The retroviral vector pRV100G backbone was used to insert TOP1MT open reading frame (ORF) followed by C-terminal T2A mCherry tag. Site directed mutagenesis was used to introduce the C578T variant in TOP1MT. The empty vector control was missing the TOP1MT ORF and only contained the backbone with T2A mCherry.

### Generation of stable lines

Stable cells expressing wild type TOP1MT (NM_052963.2), TOP1MT P193L variant, or empty vector controls were generated via retroviral transduction. Briefly, Phoenix cells were transfected with the retroviral plasmids. Then the HCT116 WT or HCT116 KO cells were transduced using 5 ml of the supernatant containing virus after 48 and 72 hrs in 100 mm plates. Following transduction, Flow cytometry sorting was done for the red fluorescence from approximately 6X10^6^ cells were using a 130 μm nozzle on a BD FACSAria Fusion (FACSAriaIII) cytometer (BD Biosciences), supported by FACSDiva Version 8.0.1. As for the cells with low expression of TOP1MT, all the positively expressing red fluorescent cells were subjected to sorting again separating them into low threshold, intermediate and high expression red fluorescence gates.

### PCR and sequencing

PureLink Genomic DNA Mini Kit (Thermo Fisher Scientific, K182001) was used for total DNA extraction. TOP1MT cDNA was amplified by PCR using the forward primer CACAACAAAGGAGGTTTTCCGGAAG and the reverse primer TGCAGTCCTTCCCCAGGA to confirm the success of the transduction. The amplified band was further purified and sequenced using Sanger sequencing to confirm the variant status of TOP1MT.

### Live cell imaging

Live cell imaging was done on 35 mm glass bottom dishes, 8L×L10^4^ cells were seeded and grown for two days. MitoTracker Red (50 nM) (Thermo Fisher Scientific, M7512) and PicoGreen (Thermo Fisher Scientific, P7581) (3 μL/mL) were used to visualize mitochondrial networks and mtDNA nucleoids respectively(39, 93).

### Immunofluorescence staining

1.5L×L10^4^ cells were seeded over 12 mm glass coverslips (no. 1.5) in 24 wells plates and incubated for 2 days. Subsequently, cells were fixed with 4% paraformaldehyde and stained with mitochondrial networks labeled with a primary antibody against TOMM20 (Sigma-Aldrich, HPA011562) used at 1:1000, STING (D2P2F) (Cell signaling, 13647S) used at 1:300, endoplasmic reticulum ER Calnexin (Millipore Sigma, MAB3126) at 1:1000 and Golgi TGN46 (Proteintech, 66477-1-Ig) used at 1:500. The corresponding Alexa fluor-conjugated secondary antibodies (Thermo Fisher Scientific) were used at 1:1000.

### Fluorescence in situ hybridization (FISH)

Cells grown on fibronectin-coated coverslips (Fisher Scientific #12-545-81) were fixed at 37°C using a solution of 4% paraformaldehyde in PHEM buffer (60 mM PIPES, 25 mM HEPES, 10 mM EGTA, 4 mM MgSO4, pH 6.8) for 15 minutes, and then permeabilized with 0.1% (v/v) Triton X-100 in PBS for 10 minutes at room temperature. FISH was then performed as previously described(94, 95), using the mREP probe conjugated to Atto633, which was purchased from Integrated DNA Technologies. For immunofluorescence, coverslips were blocked with filtered PBS containing 1% (w/v) BSA at room temperature for at least an hour, following either permeabilization or FISH. Incubation with primary antibodies was carried out in PBS containing 1% (w/v) BSA at 4°C overnight, followed by 4 x 5 minute washes in PBS. Secondary antibodies were incubated in PBS containing 1% BSA for 1 hour at room temperature. Secondary antibody was removed by 4 x 5 minute washes in PBS. Coverslips were then mounted onto slides using Prolong Glass. The following antibodies were obtained commercially and used at the indicated dilutions for immunofluorescence: cGAS (CST #15102, 1:100), HSP60 (EnCor Biotechnology #CPCA-HSP60, 1:500), DNA (Millipore Sigma #CBL186, 1:200). Secondary antibodies conjugated to Alexa 405, 488 or 647 were purchased from Thermo Fisher and used at 1:500, with the exception of anti-chicken Alexa 405, which was from Abcam and used at 1:250.

### Microscopy

An Olympus spinning disc confocal system (Olympus SD OSR) (UAPON 100XOTIRF/1.49 oil objective) was used to image fixed samples and the microscope was operated by Metamorph software. Whenever there was live imaging, a cellVivo incubation module was used to maintain cells at 37 °C and 5% CO_2_. For FISH and nucleoid-cGAS overlap images, imaging was performed using a Plan-Apochromat ×63/1.4 NA oil objective on an inverted Zeiss spinning disk microscope using 405, 488, 561, and 633 nm laser lines.

### Image analysis for mtDNA nucleoids

The particle analysis tool in ImageJ FIJI was used to measure nucleoid size after binarizing the images(96). A region of interest in cells stained with PicoGreen was manually selected including all the mitochondrial network of the cell but excluding the nucleus, then the surface area of each nucleoid was automatically determined using the analyze particles feature. The nucleoids from at least 10 cells were measured. Data was represented by a violin plot representing the distribution of the average mtDNA nucleoid sizes per cell for the indicated cell lines. P values were based on unpaired, 2-tailed Student’s t-tests.

### Image analysis for mitochondrial networks

Images of the cells stained with anti-Tomm20 (Sigma-Aldrich, HPA011562) were used for qualitative evaluation of the mitochondrial network morphology. The network was classified into three categories, fragmented (predominantly small mitochondrial fragments), intermediate (cells containing a mixture of short fragments and elongated networks) and fused (elongated, interconnected networks with few to no short fragments)(93). At least 50 cells from three independent biological replicates were scored for each of the investigated cell lines. The results shown represent meansL±LSEM, and P values were based on unpaired, 2-tailed Student’s t- tests.

### Image analysis for STING localization

Images of the cells stained with anti-STING (D2P2F) (Cell signaling, 13647S), anti-Calnexin (Millipore Sigma, MAB3126) and anti-TGN46 (Proteintech, 66477-1-Ig) were used for quantitative evaluation of the localization of STING to either the ER or the Golgi. Images were binarized, then the surface area of the total STING signal or the localized signal to the corresponding organelle were measured. The ratio of the localized area to the total STING area was determined from at least 10 images. Data was represented as meansL±LSEM, and P values were based on unpaired, 2-tailed Student’s t-tests.

### Image analysis of cytoplasmic nucleoids and overlap with cGAS

The mitochondria channel was masked using automatic thresholding, and this mask was subtracted from the mtDNA FISH channel using the image calculator in Fiji. Any remaining mtDNA FISH signal that did not overlap with anti-DNA immunofluorescent puncta were subtracted from analysis. The number of remaining mtDNA FISH puncta, as well as their overlap with cGAS, was scored.

### mtDNA copy number analysis

The PureLink Genomic DNA Mini Kit (Thermo Fisher Scientific, K182001) was used to isolate total DNA from cultured cells using according to the manufacturer’s protocol. Quantitative PCR (QPCR) was used to determine mtDNA copy number (97). Primers shown in Table 1 for mtDNA or 18S DNA were used with PowerUp SYBR Green Master Mix (Thermo Fisher Scientific, A25742) to amplify 50-100 ng of DNA template, in the QuantStudio 6 Flex Real-Time PCR system (Thermo Fisher Scientific) machine. The delta delta Ct method was used to determine mtDNA copy number using the 18S gene as a reference. Reactions were performed in three technical replicates and mtDNA copy number analysis was performed on at least three independent biological replicates. Data is presented as meanL±LSEM and unpaired, 2-tailed Student’s t-tests were used to determine statistical significance.

**Table 1:**
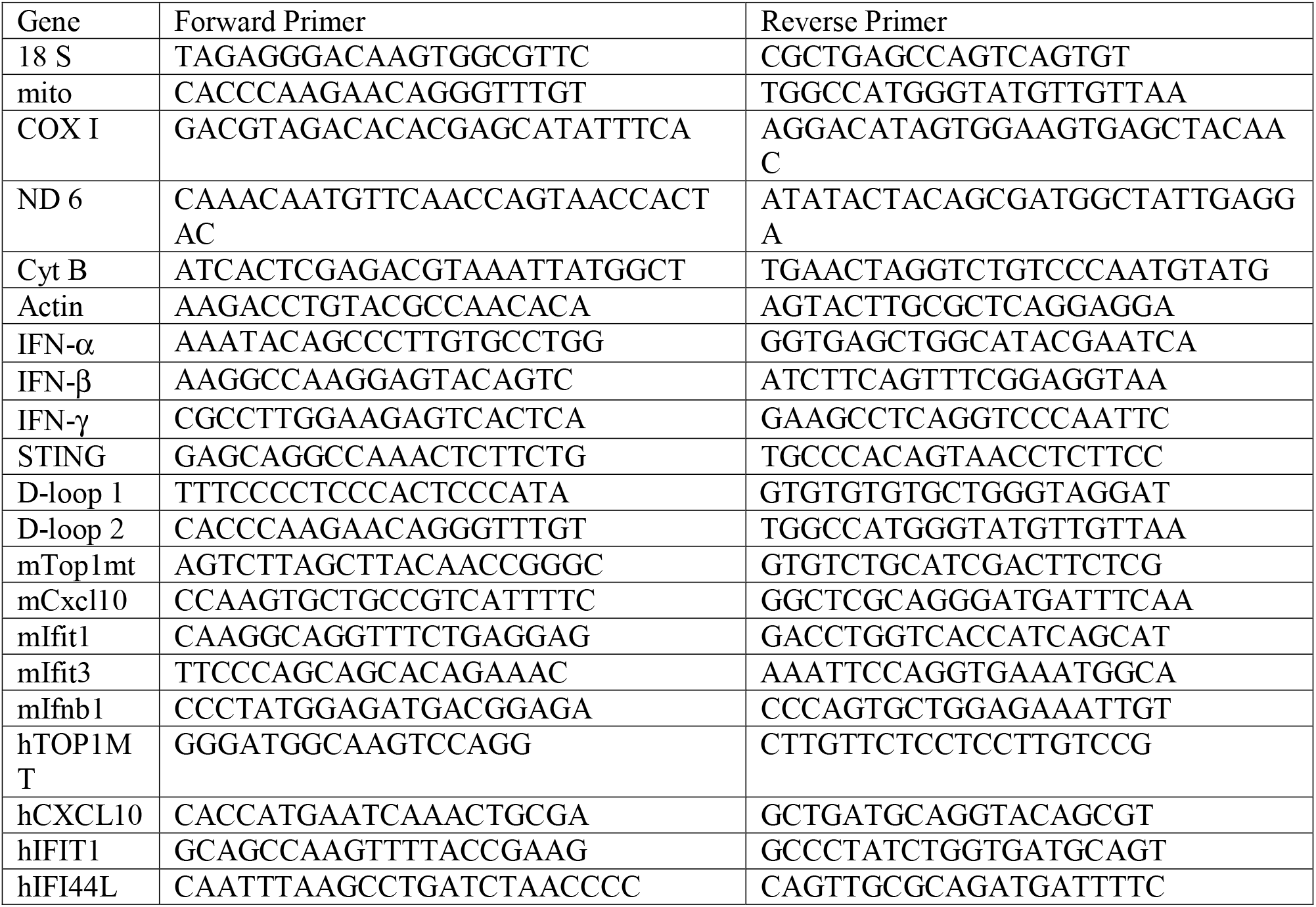
Primers used for quantitative PCR for detection of indicated genes.

### Ethidium bromide depletion/repletion

1μg/ml Ethidium bromide (Fisher Scientific, 15-585-011) was used to deplete the cells from their mtDNA content for three days, as described previously(54), then fresh media was. added afterwards to allow the cells to recover their mtDNA copy number. Cell pellets from the days 0, 3, 6 and 8 were collected for DNA purification and copy number analysis.

### Western blot

RIPA buffer (Thermo Scientific™, 89900) containing protease inhibitors was used to lyse collected cell pellets after washing with 1× phosphate buffered saline (PBS). SDS-PAGE gels were used to resolve 50 μg of total cell lysates from different cell lines then PVDF membranes were used for the transferred blot. Blots were probed with several antibodies at the indicated dilutions; OxPhos antibody cocktail (Abcam, ab110411; 1:500 dilution), anti-Actin (Sigma, A5316; 1:1000), anti-MTCO2 (Abcam, ab110258; 1:1000), anti-GAPDH (Millipore Sigma, ABS16; 1:1000), anti-TOP1MT-3 (Developmental Studies Hybridoma Bank, CPTC- TOP1MT-3; 1:200). Appropriate horseradish peroxidase (HRP)-conjugated secondary antibodies were used at 1:3000 as follows: goat anti rabbit IgG, HRP linked Antibody (Cell Signaling Technology, 7074S), or goat anti-mouse IgG-HRP (Santa Cruz Biotechnology, sc-2055). The Clarity ECL substrate (Biorad, 1705061) was used to expose the horseradish enzyme attached to each antibody using an Amersham Imager AI600.

### Southwestern blot

Cells were treated with 10 uM Aphidicolin (HelloBio Ltd, Bristol UK) for 4 hours to inhibit nuclear DNA synthesis, then 50 uM BrdU (Invitrogen, fisher scientific) is added for another 24 hours. Total DNA was isolated from 2X10^7^ cells according to the instruction of E.Z.N.A Tissue DNA kit. As controls, 1ug of DNA was treated with 10U of Topoisomerase I (New England Biolabs) at 37°C for 1 hour, or BamHI 20 U (New England Biolabs) at 37°C for 1 hour. DNA samples (1ug) were then separated on a 0.45% agarose gel (Biotechnology VWR, life science) at 30V 18-20 hours in cold room. The gel was washed briefly with water and soaked in denaturing buffer (0.5N NaOH, 1.5M NaCl) for 45-60 mins. The gel was washed again in water and soaked in 10X saline-sodium citrate (SSC) for 30 mins. Transfer to PVDF membrane (Bio-Rad, Cat# 1620177) was done using capillary action as described previously (Nature protocols, 518, 2006). The PVDF membrane was washed in 6X SSC for two minutes, and the DNA bands on the membrane were immobilized by UV-crosslinking (UV Stratalinker 1800). After blocking with 10% non-fat dry milk for 1h, the PVDF membrane was incubated for 2 h at 20 °C with anti-BrdU antibody (1:2000X dilution in TBST, Becton Dickinson Immunocytometry Systems), followed by incubations with corresponding horseradish peroxidase (HRP)-conjugated goat polyclonal anti-mouse IgG (1:5,000 dilution, EMD Millipore Etobicoke, Ontario, Canada). Blots were developed with Supersignal West Femto enhanced chemiluminescence substrate (Thermo Scientific) and imaged using the chemiluminescence imaging analyzer (LAS3000mini; Fujifilm, Tokyo, Japan).

### RNA extraction and quantification

The E.Z.N.A.® HP Total RNA Kit, Omega Bio-tek® (VWR Scientific, CA101414-850) was used for RNA extraction from cell pellets according to the manufacturer’s protocol.

Following extraction, 7.5 μg of RNA was reverse transcribed by iScript™ Advanced cDNA Synthesis Kit (Biorad, 1725038) to generate cDNA. Quantitative PCR with PowerUp SYBR Green Master Mix (Thermo Fisher Scientific, A25742) was used to amplify the genes of interest according to the primers mentioned in the Table 1 from 15-20 ng of cDNA in the QuantStudio 6 Flex Real-Time PCR system (Thermo Fisher Scientific) machine. Actin expression was used as the control in the delta delta Ct method to determine the relative expression of various transcripts. Reactions were performed in triplicate and mtDNA copy number analysis was performed on at least three independent biological replicates. Data is presented as meanL±LSEM and unpaired, 2-tailed Student’s t-tests were used to determine statistical significance.

### Cytosolic mtDNA quantification

Cell pellet was resuspended in roughly 500 µl buffer containing 150 mM NaCl, 50 mM HEPES, pH 7.4, and 15–25 µg/ml digitonin (Sigma Aldrich, D141). The homogenates were incubated end over end for 25 minutes to allow selective plasma membrane permeabilization, then centrifuged at 980 g for 10 min to pellet intact cells. The first pellet was used to extract total DNA using the PureLink Genomic DNA Mini Kit (Thermo Fisher Scientific, K182001). The cytosolic supernatants were transferred to fresh tubes and spun at 17000 g for 10 min to pellet any remaining cellular debris, yielding cytosolic preps free of nuclear, mitochondrial, and ER contamination. DNA was then isolated from these pure cytosolic fractions using the PureLink Genomic DNA Mini Kit (Thermo Fisher Scientific, K182001). qPCR was performed on both whole cell extracts and cytosolic fractions using nuclear DNA primers (18S) and mtDNA primers (D loop1-2, CytB, COX I, ND 6) as indicated in Table 1, and the CT values obtained for mtDNA abundance for whole cell extracts served as normalization controls for the mtDNA values obtained from the cytosolic fractions. This allowed effective standardization among samples and controlled for any variations in the total amount of mtDNA in control and stable cell lines samples.

### cGAS inhibition and recovery

RU.521 (Invivogen, inh-ru521) was used at a concentration of 5 ug/ml to inhibit cGAS in cells seeded in 10 cm plates, after 24 hours fresh media was added to the cells to recover for another 24 hours. Similar volume of DMSO was used as a negative control to eliminate any effect of the drug solvent.

### Structural modeling

Structural models for the TOP1MT variants were generated via RoseTTAFold, a deep learning solution to predicting protein structures, which was conducted on the Robetta web server(98). Five structures were generated per mutant, the structure with the highest confidence level was selected for further use in analysis. Once the structures were selected for each mutant, HADDOCK was used to perform docking(99, 100). The binding strength and location of the binding were approximated to generate models with both short (10) and long (50) strands of mtDNA, with sequences detailed in the following reference(36, 90). Short strand docking offered insight into the highly charged barrel through the middle of the protein. Long strand modelling then mixed the barrel analysis with the stabilizing forces of the surrounding protein structure.

The most probable binding sites were identified and fit by HADDOCK and the model selected was the one with the best HADDOCK score. These models and the interaction lists generated around them were used then to compare the binding efficacy and form between the mutants and the wild type. Models were downloaded, screened and visualized in Pymol(101).

### TCOP1MT Knockdown

siRNA transfection was performed with 25 nM siRNA duplexes (Integrated DNA Techonologies) shown in the table 2 and 3 uL of Lipofectamine RNAiMAX reagent (ThermoFisher Scientific, 13778150) according to the manufacturer’s protocol.

**Table 2:** siRNA duplexes sequences used to knockdown TOP1MT in MEF and HFF-1 cells.

|  | Sense | antisense |
| --- | --- | --- |
| Mouse<br>siTop1<br>mt | rCrCrUrGrCrArArArGrUrCrUrUrArGrCr<br>UrUrArCrArArCCG | rCrGrGrUrUrGrUrArArGrCrUrArArGrArCrUr<br>UrUrGrCrArGrGrUrU |
| Human<br>siTOP1<br>MT | rCrArGrCrUrArArGrArUrCrUrUrArUrCr<br>CrUrArCrArArCCG | rCrGrGrUrUrGrUrArGrGrArUrArArGrArUrCr<br>UrUrArGrCrUrGrCrU |

**Table 3.** Summary of analyzed phenotypes for high levels of P193L variant. Analyzed phenotypes where +++ represents full rescue, ++ represent moderate rescue, + for weak rescue and – for no rescue at all.

|  | **KO-ctrl** | **Rescue** | **P193L** |
| --- | --- | --- | --- |
| Nucleoid size | Smaller | +++ | + |
| Supercoiling | Higher | +++ | ++ |
| mtDNA Copy number | Higher | +++ | +++ |
| mtDNA Repletion | Slower | +++ | ++ |
| Mitochondrial Transcripts | Lower | +++ | + |
| OXPHOS Proteins | Lower | +++ | + |
| Mitochondrial Oxygen consumption | Lower | +++ | - |
| Mitochondrial morphology | Fused | +++ | + |
| cGAS-STING activation | Activated | +++ | ++ |

## Acknowledgement

**Funding:** This work was supported by funds provided by the Alberta Children’s Hospital Research Institute (ACHRI), the Alberta Children’s Hospital Foundation (ACHF) and the Natural Sciences and Engineering Research Council of Canada (NSERC - RGPIN-2016-04083) (T.S.). W.T.G. was supported by a British Columbia Children’s Hospital Institute (BCCHR) Intramural Investigator Grant Award Program (IGAP) award, and by a catalyst grant from Lupus Canada. Also, this work was supported by National Institutes of Health grants K99GM141482 to L.E.N. and R01 AR069876 to G.S.S., who also holds the Audrey Geisel Chair in Biomedical Science. This work was also supported by the NIH Intramural Program, the Center for Cancer Research, US National Cancer Institute (BC006161 to Y.P., H.Z and S.H.). A.K. received research funding from MitoCanada. Y.L., S.T-O., and A.P.W. were supported by NIH grant R01HL148153. S.T.-O. was supported by a Ruth L. Kirschstein National Research Service Award (NRSA) Predoctoral Fellowship to Promote Diversity in Health-Related Research F31HL160141. I.A.K. is a recipient of an ACHRI Graduate Studentship and William H. Davies scholarship. The funding sources were not involved in the study design, data collection and analysis, the writing of the report, nor in the decision to submit.

## Author contributions

I.A.K. conceived the study, designed, performed, and analyzed experiments and wrote the manuscript. J.D. performed southwestern blot and cytosolic mtDNA experiments. Y.L and S. T.-O. performed the work in mouse and human fibroblast lines. G.R.R. and L.E.N. performed the FISH experiments. B.K.C. performed the experiments on patient-derived lymphoblastoid cell lines. A.S. generated the structural model. H.Z., S.N.H. and Y.P. generated the HCT116 *TOP1MT* KO cells, shared the TOP1MT cDNA, and provided input on the project. A.K. provided input on the project and edited the manuscript. G.S.S. and A.P.W. helped design and supervise the study. W.T.G. identified the family bearing the *TOP1MT* patient variants, including clinical and family history, provided input on the project, and edited the manuscript. T.S. conceived and designed the study, analyzed experiments, supervised the study, and wrote the manuscript.

## References

1 Wincup, C. and Radziszewska, A. (2021) Abnormal Mitochondrial Physiology in the Pathogenesis of Systemic Lupus Erythematosus. Rheum Dis Clin North Am, 47, 427–439.

2 Yang, S.K., Zhang, H.R., Shi, S.P., Zhu, Y.Q., Song, N., Dai, Q., Zhang, W., Gui, M. and Zhang, H. (2020) The Role of Mitochondria in Systemic Lupus Erythematosus: A Glimpse of Various Pathogenetic Mechanisms. Curr Med Chem, 27, 3346–3361.

3 Alperin, J.M., Ortiz-Fernández, L. and Sawalha, A.H. (2018) Monogenic Lupus: A Developing Paradigm of Disease. Front Immunol, 9, 2496.

4 Bursztejn, A.C., Briggs, T.A., del Toro Duany, Y., Anderson, B.H., O’Sullivan, J., Williams, S.G., Bodemer, C., Fraitag, S., Gebhard, F., Leheup, B. et al. (2015) Unusual cutaneous features associated with a heterozygous gain-of-function mutation in IFIH1: overlap between Aicardi-Goutières and Singleton-Merten syndromes. Br J Dermatol, 173, 1505–1513.

5 An, J., Briggs, T.A., Dumax-Vorzet, A., Alarcón-Riquelme, M.E., Belot, A., Beresford, M., Bruce, I.N., Carvalho, C., Chaperot, L., Frostegård, J. et al. (2017) Tartrate-Resistant Acid Phosphatase Deficiency in the Predisposition to Systemic Lupus Erythematosus. Arthritis Rheumatol, 69, 131–142.

6 Crow, Y.J., Chase, D.S., Lowenstein Schmidt, J., Szynkiewicz, M., Forte, G.M., Gornall, H.L., Oojageer, A., Anderson, B., Pizzino, A., Helman, G. et al. (2015) Characterization of human disease phenotypes associated with mutations in TREX1, RNASEH2A, RNASEH2B, RNASEH2C, SAMHD1, ADAR, and IFIH1. Am J Med Genet A, 167a, 296–312.

7 Dyall, S.D., Brown, M.T. and Johnson, P.J. (2004) Ancient invasions: from endosymbionts to organelles. Science (New York, N.Y.), 304, 253–257.

8 Kukat, C., Wurm, C.A., Spahr, H., Falkenberg, M., Larsson, N.G. and Jakobs, S. (2011) Super-resolution microscopy reveals that mammalian mitochondrial nucleoids have a uniform size and frequently contain a single copy of mtDNA. Proceedings of the National Academy of Sciences of the United States of America, 108, 13534–13539.

9 Kausar, S., Yang, L., Abbas, M.N., Hu, X., Zhao, Y., Zhu, Y. and Cui, H. (2020) Mitochondrial DNA: A Key Regulator of Anti-Microbial Innate Immunity. Genes (Basel), 11.

10 Cheng, Z., Abrams, S.T., Austin, J., Toh, J., Wang, S.S., Wang, Z., Yu, Q., Yu, W., Toh, C.H. and Wang, G. (2020) The Central Role and Possible Mechanisms of Bacterial DNAs in Sepsis Development. Mediators Inflamm, 2020, 7418342.

11 Luna-Sánchez, M., Bianchi, P. and Quintana, A. (2021) Mitochondria-Induced Immune Response as a Trigger for Neurodegeneration: A Pathogen from Within. Int J Mol Sci, 22.

12. Riley, J.S. and Tait, S.W. (2020) Mitochondrial DNA in inflammation and immunity. EMBO reports, 21, e49799.

13 Lepelley, A., Wai, T. and Crow, Y.J. (2021) Mitochondrial Nucleic Acid as a Driver of Pathogenic Type I Interferon Induction in Mendelian Disease. Front Immunol, 12, 729763.

14 Yamazaki, T. and Galluzzi, L. (2022) BAX and BAK dynamics control mitochondrial DNA release during apoptosis. Cell Death Differ, 29, 1296–1298.

15 Patrushev, M., Kasymov, V., Patrusheva, V., Ushakova, T., Gogvadze, V. and Gaziev, A. (2004) Mitochondrial permeability transition triggers the release of mtDNA fragments. Cell Mol Life Sci, 61, 3100–3103.

16 Kim, J., Gupta, R., Blanco, L.P., Yang, S., Shteinfer-Kuzmine, A., Wang, K., Zhu, J., Yoon, H.E., Wang, X., Kerkhofs, M. et al. (2019) VDAC oligomers form mitochondrial pores to release mtDNA fragments and promote lupus-like disease. Science (New York, N.Y.), 366, 1531–1536.

17 Fang, C., Wei, X. and Wei, Y. (2016) Mitochondrial DNA in the regulation of innate immune responses. Protein Cell, 7, 11–16.

18 Guo, Y., Gu, R., Gan, D., Hu, F., Li, G. and Xu, G. (2020) Mitochondrial DNA drives noncanonical inflammation activation via cGAS-STING signaling pathway in retinal microvascular endothelial cells. Cell Commun Signal, 18, 172.

19 Huang, L.S., Hong, Z., Wu, W., Xiong, S., Zhong, M., Gao, X., Rehman, J. and Malik, A.B. (2020) mtDNA Activates cGAS Signaling and Suppresses the YAP-Mediated Endothelial Cell Proliferation Program to Promote Inflammatory Injury. Immunity, 52, 475–486.e475.

20 Bai, J., Cervantes, C., He, S., He, J., Plasko, G.R., Wen, J., Li, Z., Yin, D., Zhang, C., Liu, M. et al. (2020) Mitochondrial stress-activated cGAS-STING pathway inhibits thermogenic program and contributes to overnutrition-induced obesity in mice. Commun Biol, 3, 257.

21 Decout, A., Katz, J.D., Venkatraman, S. and Ablasser, A. (2021) The cGAS-STING pathway as a therapeutic target in inflammatory diseases. Nat Rev Immunol, 21, 548–569.

22 Dobbs, N., Burnaevskiy, N., Chen, D., Gonugunta, V.K., Alto, N.M. and Yan, N. (2015) STING Activation by Translocation from the ER Is Associated with Infection and Autoinflammatory Disease. Cell Host Microbe, 18, 157–168.

23 West, A.P., Khoury-Hanold, W., Staron, M., Tal, M.C., Pineda, C.M., Lang, S.M., Bestwick, M., Duguay, B.A., Raimundo, N., MacDuff, D.A. et al. (2015) Mitochondrial DNA stress primes the antiviral innate immune response. Nature, 520, 553–557.

24 Torres-Odio, S., Lei, Y., Gispert, S., Maletzko, A., Key, J., Menissy, S.S., Wittig, I., Auburger, G. and West, A.P. (2021) Loss of Mitochondrial Protease CLPP Activates Type I IFN Responses through the Mitochondrial DNA-cGAS-STING Signaling Axis. J Immunol, 206, 1890–1900.

25 Matsui, H., Ito, J., Matsui, N., Uechi, T., Onodera, O. and Kakita, A. (2021) Cytosolic dsDNA of mitochondrial origin induces cytotoxicity and neurodegeneration in cellular and zebrafish models of Parkinson’s disease. Nature communications, 12, 3101.

26 Sliter, D.A., Martinez, J., Hao, L., Chen, X., Sun, N., Fischer, T.D., Burman, J.L., Li, Y., Zhang, Z., Narendra, D.P. et al. (2018) Parkin and PINK1 mitigate STING-induced inflammation. Nature, 561, 258–262.

27 Du, S., Chen, G., Yuan, B., Hu, Y., Yang, P., Chen, Y., Zhao, Q., Zhou, J., Fan, J. and Zeng, Z. (2021) DNA sensing and associated type 1 interferon signaling contributes to progression of radiation-induced liver injury. Cell Mol Immunol, 18, 1718–1728.

28 Luo, X., Li, H., Ma, L., Zhou, J., Guo, X., Woo, S.L., Pei, Y., Knight, L.R., Deveau, M., Chen, Y. et al. (2018) Expression of STING Is Increased in Liver Tissues From Patients With NAFLD and Promotes Macrophage-Mediated Hepatic Inflammation and Fibrosis in Mice. Gastroenterology, 155, 1971–1984.e1974.

29 Thomsen, M.K., Skouboe, M.K., Boularan, C., Vernejoul, F., Lioux, T., Leknes, S.L., Berthelsen, M.F., Riedel, M., Cai, H., Joseph, J.V. et al. (2020) The cGAS-STING pathway is a therapeutic target in a preclinical model of hepatocellular carcinoma. Oncogene, 39, 1652–1664.

30 Maekawa, H., Inoue, T., Ouchi, H., Jao, T.M., Inoue, R., Nishi, H., Fujii, R., Ishidate, F., Tanaka, T., Tanaka, Y. et al. (2019) Mitochondrial Damage Causes Inflammation via cGAS-STING Signaling in Acute Kidney Injury. Cell reports, 29, 1261–1273.e1266.

31 Chung, K.W., Dhillon, P., Huang, S., Sheng, X., Shrestha, R., Qiu, C., Kaufman, B.A., Park, J., Pei, L., Baur, J. et al. (2019) Mitochondrial Damage and Activation of the STING Pathway Lead to Renal Inflammation and Fibrosis. Cell metabolism, 30, 784–799.e785.

32 Zhao, Q., Wei, Y., Pandol, S.J., Li, L. and Habtezion, A. (2018) STING Signaling Promotes Inflammation in Experimental Acute Pancreatitis. Gastroenterology, 154, 1822–1835.e1822.

33 Sobek, S. and Boege, F. (2014) DNA topoisomerases in mtDNA maintenance and ageing. Experimental gerontology, 56, 135–141.

34 Zhang, H., Barcelo, J.M., Lee, B., Kohlhagen, G., Zimonjic, D.B., Popescu, N.C. and Pommier, Y. (2001) Human mitochondrial topoisomerase I. Proceedings of the National Academy of Sciences of the United States of America, 98, 10608–10613.

35 Zhang, X., Li, X.H., Ma, X., Wang, Z.H., Lu, S. and Guo, Y.L. (2006) Redox-induced apoptosis of human oocytes in resting follicles in vitro. J Soc Gynecol Investig, 13, 451–458.

36 Zhang, H. and Pommier, Y. (2008) Mitochondrial topoisomerase I sites in the regulatory D-loop region of mitochondrial DNA. Biochemistry, 47, 11196–11203.

37 Zhang, H., Zhang, Y.W., Yasukawa, T., Dalla Rosa, I., Khiati, S. and Pommier, Y. (2014) Increased negative supercoiling of mtDNA in TOP1mt knockout mice and presence of topoisomerases IIα and IIβ in vertebrate mitochondria. Nucleic acids research, 42, 7259–7267.

38 Sobek, S., Dalla Rosa, I., Pommier, Y., Bornholz, B., Kalfalah, F., Zhang, H., Wiesner, R.J., von Kleist-Retzow, J.C., Hillebrand, F., Schaal, H. et al. (2013) Negative regulation of mitochondrial transcription by mitochondrial topoisomerase I. Nucleic acids research, 41, 9848–9857.

39 Al Khatib, I., Deng, J., Symes, A., Kerr, M., Zhang, H., Huang, S.N., Pommier, Y., Khan, A. and Shutt, T.E. (2022) Functional characterization of two variants of mitochondrial topoisomerase TOP1MT that impact regulation of the mitochondrial genome. The Journal of biological chemistry, 298, 102420.

40 Han, S., Zhuang, H., Shumyak, S., Yang, L. and Reeves, W.H. (2015) Mechanisms of autoantibody production in systemic lupus erythematosus. Front Immunol, 6, 228.

41 Yaniv, G., Twig, G., Shor, D.B., Furer, A., Sherer, Y., Mozes, O., Komisar, O., Slonimsky, E., Klang, E., Lotan, E. et al. (2015) A volcanic explosion of autoantibodies in systemic lupus erythematosus: a diversity of 180 different antibodies found in SLE patients. Autoimmun Rev, 14, 75–79.

42 Rekvig, O.P. (2015) Anti-dsDNA antibodies as a classification criterion and a diagnostic marker for systemic lupus erythematosus: critical remarks. Clin Exp Immunol, 179, 5–10.

43 Bai, Y., Tong, Y., Liu, Y. and Hu, H. (2018) Self-dsDNA in the pathogenesis of systemic lupus erythematosus. Clin Exp Immunol, 191, 1–10.

44 Fredi, M., Cavazzana, I., Zanola, A., Carabellese, N., Tincani, A., Mahler, M. and Franceschini, F. (2017) Anti-topoisomerase-I antibodies in systemic lupus erythematosus and potential association with the presence of anti-dsDNA antibodies. Lupus, 26, 1121–1122.

45 Stojanov, L., Satoh, M., Dooley, M.A., Kuwana, M., Jennette, J.C. and Reeves, W.H. (1995) Autoantibodies to topoisomerase I in a patient with systemic lupus erythematosus without features of scleroderma. Lupus, 4, 314–317.

46 Hamidou, M.A., Audrain, M.A., Masseau, A., Agard, C. and Moreau, A. (2006) Anti-topoisomerase I antibodies in systemic lupus erythematosus as a marker of severe nephritis. Clin Rheumatol, 25, 542–543.

47 Mahler, M., Silverman, E.D., Schulte-Pelkum, J. and Fritzler, M.J. (2010) Anti-Scl-70 (topo-I) antibodies in SLE: Myth or reality? Autoimmun Rev, 9, 756–760.

48 Budde, P., Zucht, H.D., Vordenbäumen, S., Goehler, H., Fischer-Betz, R., Gamer, M., Marquart, K., Rengers, P., Richter, J., Lueking, A. et al. (2016) Multiparametric detection of autoantibodies in systemic lupus erythematosus. Lupus, 25, 812–822.

49 Thim-Uam, A., Prabakaran, T., Tansakul, M., Makjaroen, J., Wongkongkathep, P., Chantaravisoot, N., Saethang, T., Leelahavanichkul, A., Benjachat, T., Paludan, S. et al. (2020) STING Mediates Lupus via the Activation of Conventional Dendritic Cell Maturation and Plasmacytoid Dendritic Cell Differentiation. iScience, 23, 101530.

50 Li, Y., Wilson, H.L. and Kiss-Toth, E. (2017) Regulating STING in health and disease. J Inflamm (Lond), 14, 11.

51 Jeremiah, N., Neven, B., Gentili, M., Callebaut, I., Maschalidi, S., Stolzenberg, M.C., Goudin, N., Frémond, M.L., Nitschke, P., Molina, T.J. et al. (2014) Inherited STING-activating mutation underlies a familial inflammatory syndrome with lupus-like manifestations. J Clin Invest, 124, 5516–5520.

52 Arneth, B. (2019) Systemic Lupus Erythematosus and DNA Degradation and Elimination Defects. Front Immunol, 10, 1697.

53 Gibson, W.T., Farooqi, I.S., Moreau, M., DePaoli, A.M., Lawrence, E., O’Rahilly, S. and Trussell, R.A. (2004) Congenital leptin deficiency due to homozygosity for the Delta133G mutation: report of another case and evaluation of response to four years of leptin therapy. J Clin Endocrinol Metab, 89, 4821–4826.

54 Khiati, S., Baechler, S.A., Factor, V.M., Zhang, H., Huang, S.Y., Dalla Rosa, I., Sourbier, C., Neckers, L., Thorgeirsson, S.S. and Pommier, Y. (2015) Lack of mitochondrial topoisomerase I (TOP1mt) impairs liver regeneration. Proceedings of the National Academy of Sciences of the United States of America, 112, 11282–11287.

55 Vincent, J., Adura, C., Gao, P., Luz, A., Lama, L., Asano, Y., Okamoto, R., Imaeda, T., Aida, J., Rothamel, K. et al. (2017) Small molecule inhibition of cGAS reduces interferon expression in primary macrophages from autoimmune mice. Nature communications, 8, 750.

56 Suter, M.A., Tan, N.Y., Thiam, C.H., Khatoo, M., MacAry, P.A., Angeli, V., Gasser, S. and Zhang, Y.L. (2021) cGAS-STING cytosolic DNA sensing pathway is suppressed by JAK2- STAT3 in tumor cells. Scientific reports, 11, 7243.

57 Pépin, G. and Gantier, M.P. (2018) Assessing the cGAS-cGAMP-STING Activity of Cancer Cells. Methods in molecular biology (Clifton, N.J.), 1725, 257–266.

58 Zhang, H., Zhang, Y.W., Yasukawa, T., Dalla Rosa, I., Khiati, S. and Pommier, Y. (2014) Increased negative supercoiling of mtDNA in TOP1mt knockout mice and presence of topoisomerases IIalpha and IIbeta in vertebrate mitochondria. Nucleic acids research, 42, 7259–7267.

59 Yaginuma, K., Kobayashi, M., Taira, M. and Koike, K. (1982) A new RNA polymerase and in vitro transcription of the origin of replication from rat mitochondrial DNA. Nucleic acids research, 10, 7531–7542.

60 Buzan, J.M. and Low, R.L. (1988) Preference of human mitochondrial RNA polymerase for superhelical templates with mitochondrial promoters. Biochemical and biophysical research communications, 152, 22–29.

61 Fukuoh, A., Ohgaki, K., Hatae, H., Kuraoka, I., Aoki, Y., Uchiumi, T., Jacobs, H.T. and Kang, D. (2009) DNA conformation-dependent activities of human mitochondrial RNA polymerase. Genes Cells, 14, 1029–1042.

62 Barat-Gueride, M., Dufresne, C. and Rickwood, D. (1989) Effect of DNA conformation on the transcription of mitochondrial DNA. European journal of biochemistry, 183, 297–302.

63 Zollo, O. and Sondheimer, N. (2017) Topological requirements of the mitochondrial heavy-strand promoters. Transcription, 8, 307–312.

64 Tang, J.X., Thompson, K., Taylor, R.W. and Oláhová, M. (2020) Mitochondrial OXPHOS Biogenesis: Co-Regulation of Protein Synthesis, Import, and Assembly Pathways. Int J Mol Sci, 21.

65 Baechler, S.A., Factor, V.M., Dalla Rosa, I., Ravji, A., Becker, D., Khiati, S., Miller Jenkins, L.M., Lang, M., Sourbier, C., Michaels, S.A. et al. (2019) The mitochondrial type IB topoisomerase drives mitochondrial translation and carcinogenesis. Nature communications, 10, 83.

66 Taguchi, T., Mukai, K., Takaya, E. and Shindo, R. (2021) STING Operation at the ER/Golgi Interface. Front Immunol, 12, 646304.

67 Khiati, S., Dalla Rosa, I., Sourbier, C., Ma, X., Rao, V.A., Neckers, L.M., Zhang, H. and Pommier, Y. (2014) Mitochondrial topoisomerase I (top1mt) is a novel limiting factor of doxorubicin cardiotoxicity. Clin Cancer Res, 20, 4873–4881.

68 Zhang, H., Seol, Y., Agama, K., Neuman, K.C. and Pommier, Y. (2017) Distribution bias and biochemical characterization of TOP1MT single nucleotide variants. Scientific reports, 7, 8614.

69 Wang, W., Shen, P., Thiyagarajan, S., Lin, S., Palm, C., Horvath, R., Klopstock, T., Cutler, D., Pique, L., Schrijver, I. et al. (2011) Identification of rare DNA variants in mitochondrial disorders with improved array-based sequencing. Nucleic acids research, 39, 44–58.

70 Lei, Y., Guerra Martinez, C., Torres-Odio, S., Bell, S.L., Birdwell, C.E., Bryant, J.D., Tong, C.W., Watson, R.O., West, L.C. and West, A.P. (2021) Elevated type I interferon responses potentiate metabolic dysfunction, inflammation, and accelerated aging in mtDNA mutator mice. Sci Adv, 7.

71 Lepelley, A., Della Mina, E., Van Nieuwenhove, E., Waumans, L., Fraitag, S., Rice, G.I., Dhir, A., Frémond, M.L., Rodero, M.P., Seabra, L. et al. (2021) Enhanced cGAS-STING-dependent interferon signaling associated with mutations in ATAD3A. J Exp Med, 218.

72 Kamen, D.L. (2014) Environmental influences on systemic lupus erythematosus expression. Rheum Dis Clin North Am, 40, 401–412, vii.

73 Perl, A. (2010) Pathogenic mechanisms in systemic lupus erythematosus. Autoimmunity, 43, 1–6.

74 Lood, C., Blanco, L.P., Purmalek, M.M., Carmona-Rivera, C., De Ravin, S.S., Smith, C.K., Malech, H.L., Ledbetter, J.A., Elkon, K.B. and Kaplan, M.J. (2016) Neutrophil extracellular traps enriched in oxidized mitochondrial DNA are interferogenic and contribute to lupus-like disease. Nat Med, 22, 146–153.

75 Leishangthem, B.D., Sharma, A. and Bhatnagar, A. (2016) Role of altered mitochondria functions in the pathogenesis of systemic lupus erythematosus. Lupus, 25, 272–281.

76 An, J., Durcan, L., Karr, R.M., Briggs, T.A., Rice, G.I., Teal, T.H., Woodward, J.J. and Elkon, K.B. (2017) Expression of Cyclic GMP-AMP Synthase in Patients With Systemic Lupus Erythematosus. Arthritis Rheumatol, 69, 800–807.

77 Kato, Y., Park, J., Takamatsu, H., Konaka, H., Aoki, W., Aburaya, S., Ueda, M., Nishide, M., Koyama, S., Hayama, Y. et al. (2018) Apoptosis-derived membrane vesicles drive the cGAS-STING pathway and enhance type I IFN production in systemic lupus erythematosus. Ann Rheum Dis, 77, 1507–1515.

78 Meas, R., Burak, M.J. and Sweasy, J.B. (2017) DNA repair and systemic lupus erythematosus. DNA Repair (Amst), 56, 174–182.

79 Sallai, K., Nagy, E., Derfalvy, B., Müzes, G. and Gergely, P. (2005) Antinucleosome antibodies and decreased deoxyribonuclease activity in sera of patients with systemic lupus erythematosus. Clin Diagn Lab Immunol, 12, 56–59.

80 Yasutomo, K., Horiuchi, T., Kagami, S., Tsukamoto, H., Hashimura, C., Urushihara, M. and Kuroda, Y. (2001) Mutation of DNASE1 in people with systemic lupus erythematosus. Nat Genet, 28, 313–314.

81 Al-Mayouf, S.M., Sunker, A., Abdwani, R., Abrawi, S.A., Almurshedi, F., Alhashmi, N., Al Sonbul, A., Sewairi, W., Qari, A., Abdallah, E. et al. (2011) Loss-of-function variant in DNASE1L3 causes a familial form of systemic lupus erythematosus. Nat Genet, 43, 1186–1188.

82 Zervou, M.I., Andreou, A., Matalliotakis, M., Spandidos, D.A., Goulielmos, G.N. and Eliopoulos, E.E. (2020) Association of the DNASE1L3 rs35677470 polymorphism with systemic lupus erythematosus, rheumatoid arthritis and systemic sclerosis: Structural biological insights. Mol Med Rep, 22, 4492–4498.

83 Lee-Kirsch, M.A., Gong, M., Chowdhury, D., Senenko, L., Engel, K., Lee, Y.A., de Silva, U., Bailey, S.L., Witte, T., Vyse, T.J. et al. (2007) Mutations in the gene encoding the 3’-5’ DNA exonuclease TREX1 are associated with systemic lupus erythematosus. Nat Genet, 39, 1065–1067.

84 Fye, J.M., Orebaugh, C.D., Coffin, S.R., Hollis, T. and Perrino, F.W. (2011) Dominant mutation of the TREX1 exonuclease gene in lupus and Aicardi-Goutieres syndrome. The Journal of biological chemistry, 286, 32373–32382.

85 López-López, L., Nieves-Plaza, M., Castro Mdel, R., Font, Y.M., Torres-Ramos, C.A., Vilá, L.M. and Ayala-Peña, S. (2014) Mitochondrial DNA damage is associated with damage accrual and disease duration in patients with systemic lupus erythematosus. Lupus, 23, 1133–1141.

86 Goffart, S., Hangas, A. and Pohjoismaki, J.L.O. (2019) Twist and Turn-Topoisomerase Functions in Mitochondrial DNA Maintenance. Int J Mol Sci, 20.

87 Douarre, C., Sourbier, C., Dalla Rosa, I., Brata Das, B., Redon, C.E., Zhang, H., Neckers, L. and Pommier, Y. (2012) Mitochondrial topoisomerase I is critical for mitochondrial integrity and cellular energy metabolism. PloS one, 7, e41094.

88 Piccinini, G., Cardellini, E., Reimer, G., Arnett, F.C. and Durban, E. (1991) An antigenic region of topoisomerase I in DNA polymerase chain reaction-generated fragments recognized by autoantibodies of scleroderma patients. Mol Immunol, 28, 333–339.

89 Hongliang Zhang, M.K., and Yves Pommier. Using mitochondrial TOP1 (TOP1mt) to probe the involvement of mitochondria in Systemic Lupus Erythematosus. in press.

90 Dalla Rosa, I., Huang, S.Y., Agama, K., Khiati, S., Zhang, H. Pommier, Y. (2014) Mapping topoisomerase sites in mitochondrial DNA with a poisonous mitochondrial topoisomerase I (Top1mt). The Journal of biological chemistry, 289, 18595–18602.

91 Wu, Z., Oeck, S., West, A.P., Mangalhara, K.C., Sainz, A.G., Newman, L.E., Zhang, X.O., Wu, L., Yan, Q., Bosenberg, M. et al. (2019) Mitochondrial DNA Stress Signalling Protects the Nuclear Genome. Nat Metab, 1, 1209–1218.

92 Jiang, M., Chen, P., Wang, L., Li, W., Chen, B., Liu, Y., Wang, H., Zhao, S., Ye, L., He, Y. et al. (2020) cGAS-STING, an important pathway in cancer immunotherapy. J Hematol Oncol, 13, 81.

93 Sabouny, R., Wong, R., Lee-Glover, L., Greenway, S.C., Sinasac, D.S., Khan, A. and Shutt, T.E. (2019) Characterization of the C584R variant in the mtDNA depletion syndrome gene FBXL4, reveals a novel role for FBXL4 as a regulator of mitochondrial fusion. Biochim Biophys Acta Mol Basis Dis, 1865, 165536.

94. Newman, L.E., Tadepalle, N., Novak, S.W., Schiavon, C.R., Rojas, G.R., Chevez, J.A., Lemersal, I., Medina, M., Rocha, S., Towers, C.G. et al. (2022) Endosomal removal and disposal of dysfunctional, immunostimulatory mitochondrial DNA. bioRxiv, in press., 2022.2010.2012.511955.

95 Chatre, L.R.M. (2015) Edeas, V.W.M. (ed.), In Mitochondrial Medicine. Springer, New York, Vol. Volume I, Probing Mitochondrial Function, pp. 133–147.

96 Donkervoort, S., Sabouny, R., Yun, P., Gauquelin, L., Chao, K.R., Hu, Y., Al Khatib, I., Töpf, A., Mohassel, P., Cummings, B.B. et al. (2019) MSTO1 mutations cause mtDNA depletion, manifesting as muscular dystrophy with cerebellar involvement. Acta Neuropathol, 138, 1013–1031.

97 Eaton, J.S., Lin, Z.P., Sartorelli, A.C., Bonawitz, N.D. and Shadel, G.S. (2007) Ataxia-telangiectasia mutated kinase regulates ribonucleotide reductase and mitochondrial homeostasis. J Clin Invest, 117, 2723–2734.

98 Baek, M., DiMaio, F., Anishchenko, I., Dauparas, J., Ovchinnikov, S., Lee, G.R., Wang, J., Cong, Q., Kinch, L.N., Schaeffer, R.D. et al. (2021) Accurate prediction of protein structures and interactions using a three-track neural network. Science (New York, N.Y.), 373, 871–876.

99 van Zundert, G.C.P., Rodrigues, J., Trellet, M., Schmitz, C., Kastritis, P.L., Karaca, E., Melquiond, A.S.J., van Dijk, M., de Vries, S.J. and Bonvin, A. (2016) The HADDOCK2.2 Web Server: User-Friendly Integrative Modeling of Biomolecular Complexes. J Mol Biol, 428, 720–725.

100 Honorato, R.V., Koukos, P.I., Jiménez-García, B., Tsaregorodtsev, A., Verlato, M., Giachetti, A., Rosato, A. and Bonvin, A. (2021) Structural Biology in the Clouds: The WeNMR-EOSC Ecosystem. Front Mol Biosci, 8, 729513.

101 Baker, N.A., Sept, D., Joseph, S., Holst, M.J. and McCammon, J.A. (2001) Electrostatics of nanosystems: application to microtubules and the ribosome. Proceedings of the National Academy of Sciences of the United States of America, 98, 10037–10041.

