## Supplemental Figures for "Activation of the cGAS-STING innate immune response in cells with deficient mitochondrial topoisomerase TOP1MT"

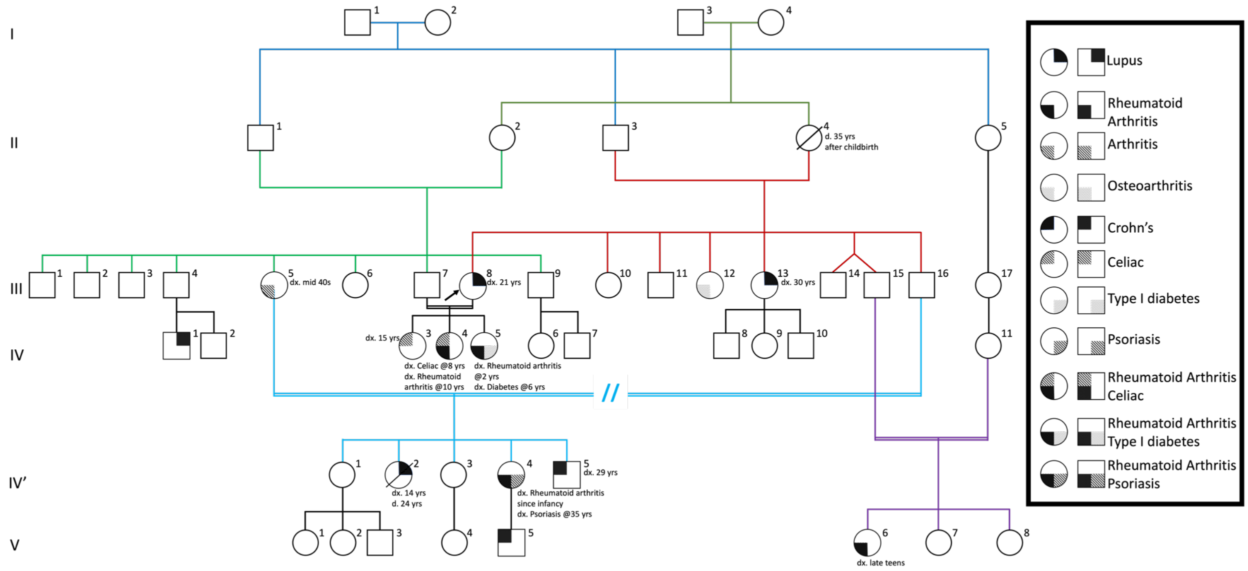


**Figure S1. Extended family pedigree.** The extended pedigree structure of the affected family shows multiple members are affected with a variety of autoimmune diseases including lupus, arthritis, Crohn’s disease, and other diseases. The legend depicts symbols used to indicate the various patient phenotypes/combination present in the pedigree. Colours are used to help visualize the different nuclear families in the larger pedigree.

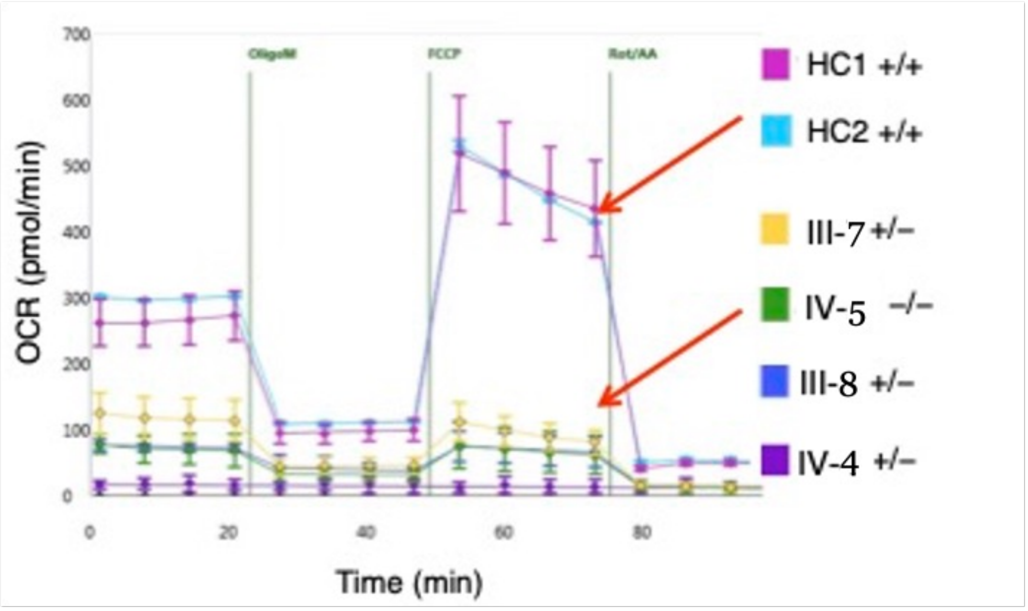


**Figure S2: Carriers of the p.(Pro193Leu) TOP1MT variant from the nuclear family of the proband have reduced oxidative phosphorylation.** Oxygen consumption rate (OCR) from lymphoblastoid cell lines was measured using a Seahorse Bioscience XF Extracellular flux analyzer for the indicated homozygous (-/-) and heterozygous (+/-) carriers of the p.(Pro193Leu) TOP1MT variant, and compared to healthy controls (HC1 and HC2, +/+).

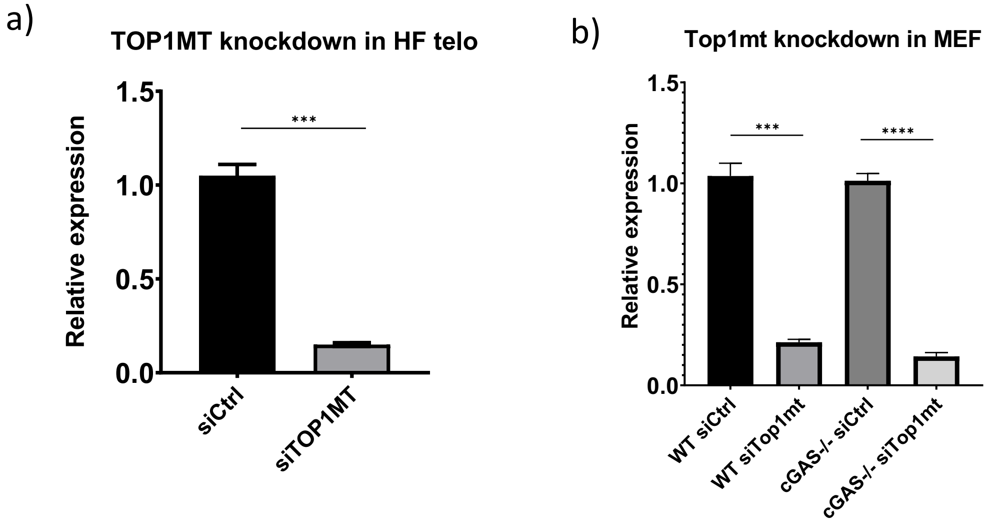


**Figure S3. Confirmation of TOP1MT siRNA knockdown.** qRT-PCR quantification of TOP1MT transcript levels in TOP1MT siRNA or control siRNA treated **(a)** human fibroblasts or **(b)** wild-type (WT) and cGAS knockout (cGAS -/-) mouse embryonic fibroblasts (MEFs). Statistical analysis was done using unpaired student t-test and p values *** <0.001 and **** <0.0001. Error bars represent standard error of mean.

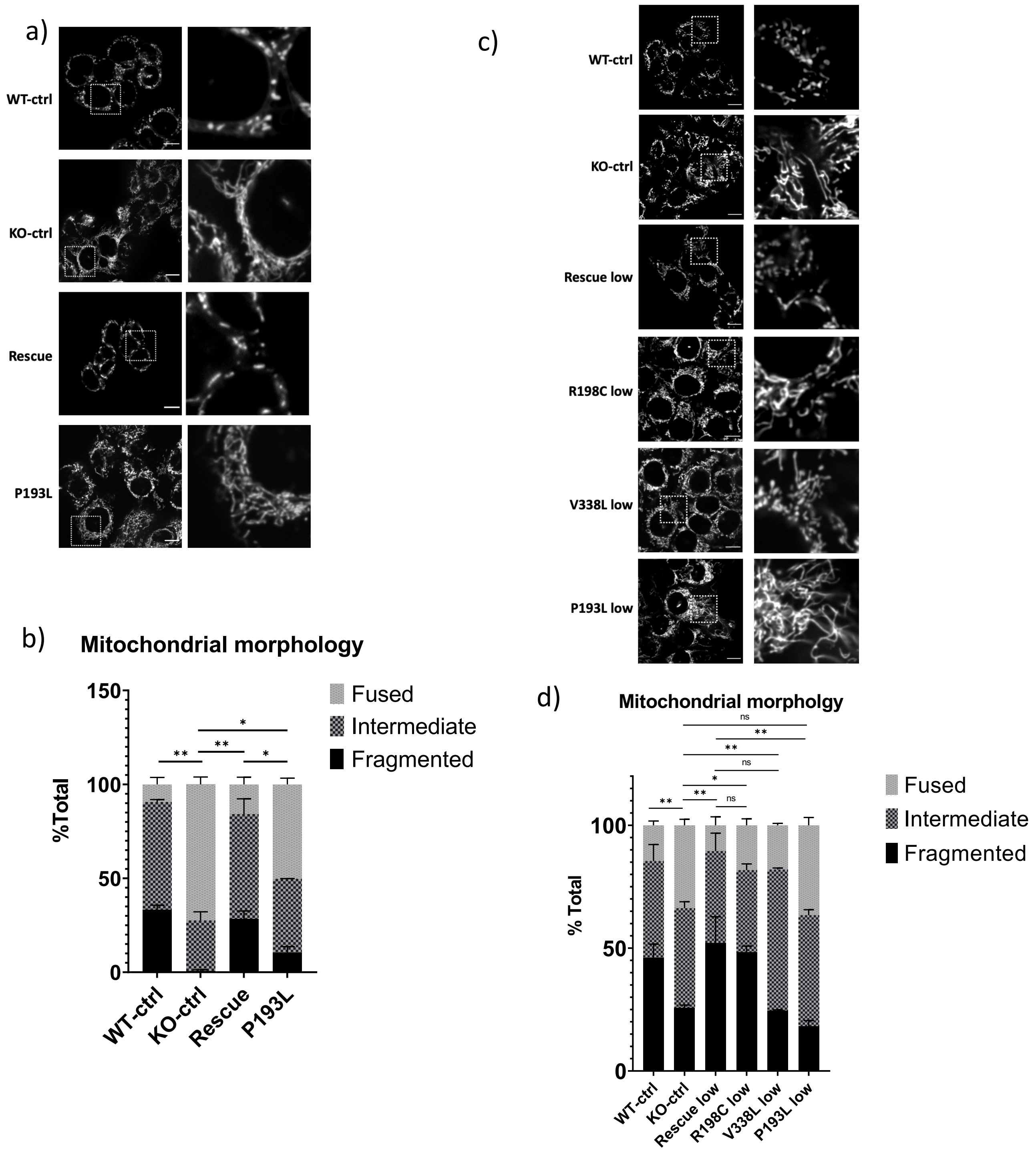


**Figure S4. TOP1MT-KO cells stably expressing P193L alter mitochondrial network. a&c)** Representative confocal images of mitochondrial networks from fixed cells for high expression (a) and low expression (c) of indicated TOP1MT variants stained with anti-TOMM20 antibody. Scalebars are 10 µm. **b&d)** Qualitative quantification of mitochondrial morphology for cells stained from (a and c). Analysis for each line was performed on three technical replicates of at least 50 cells each. All statistical analysis other than in (a) were done using unpaired student t-test and p values * <0.05 and ** <0.01 and 'ns' signifies no significant differences between indicated groups. Error bars represent standard error of mean.

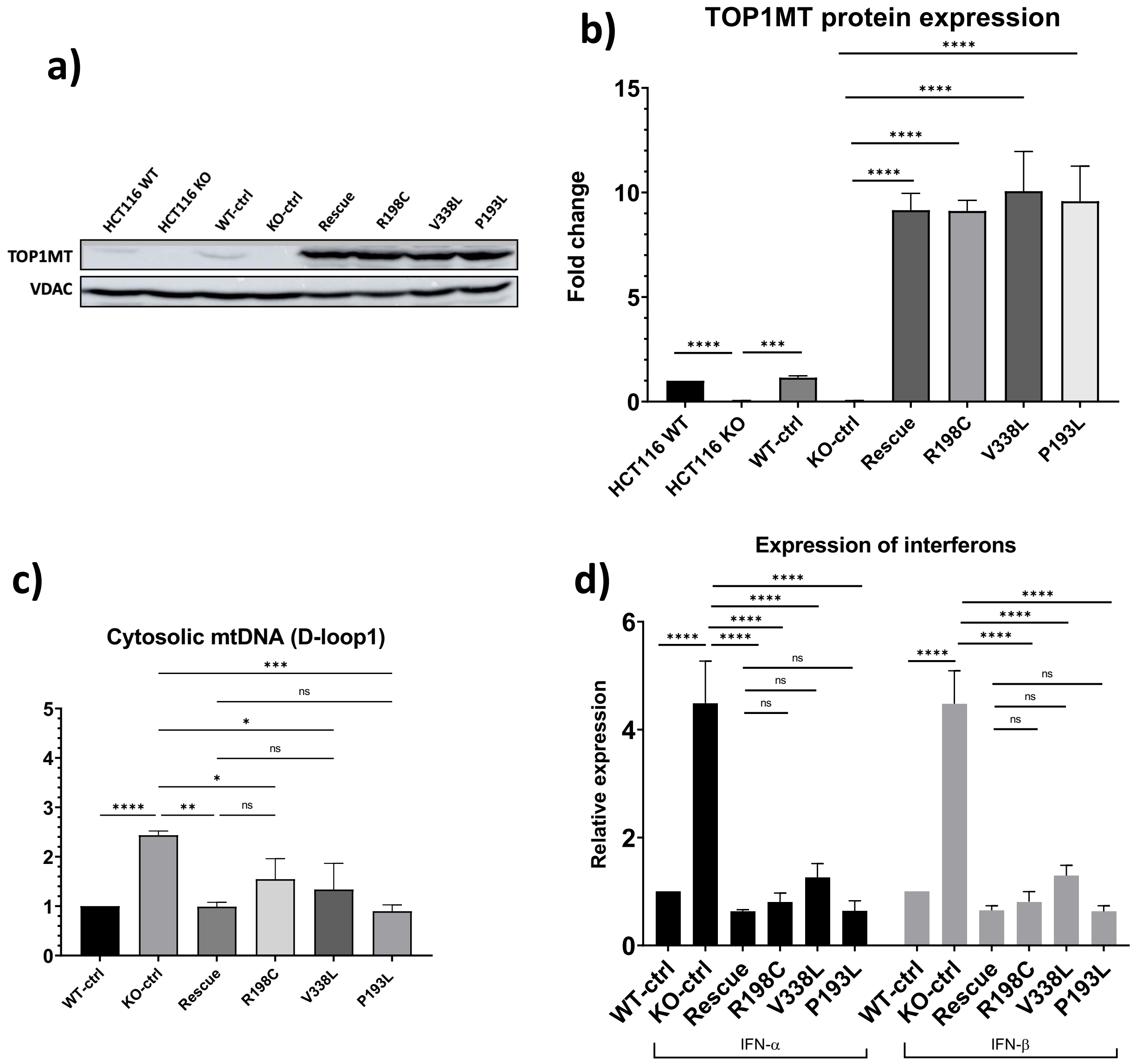


**Figure S5: Cytosolic mtDNA release and interferon levels in cell lines expressing high levels of TOP1MT. a)** Representative western blots showing elevated levels of the TOP1MT protein in cells for WT(Rescue), R198C, V338L and P193L variants. Uncropped images of western blots are available in Fig S8b. **b)** Quantification of western blots of TOP1MT protein as in (a), from three independent experiments, corrected to VDAC as a load control. **c)** qPCR quantification of cytosolic mtDNA levels from indicated cells using the primer set for the D-loop region, shows that all TOP1MT variants can reduce cytosolic mtDNA levels when expressed at high levels. Normalization was done relative to the corresponding amplicons in total DNA from the same cells. **d)** qRT-PCR analysis of type I interferon expression from indicated cells shows rescue of type I interferon levels in cells expressing high levels of all the TOP1MT variants. All statistical analysis were done using unpaired student t-test. P-values *<0.05, ** <0.01, *** <0.001 and **** <0.0001. 'ns' signifies no significant differences between indicated groups. Error bars represent standard error of mean.

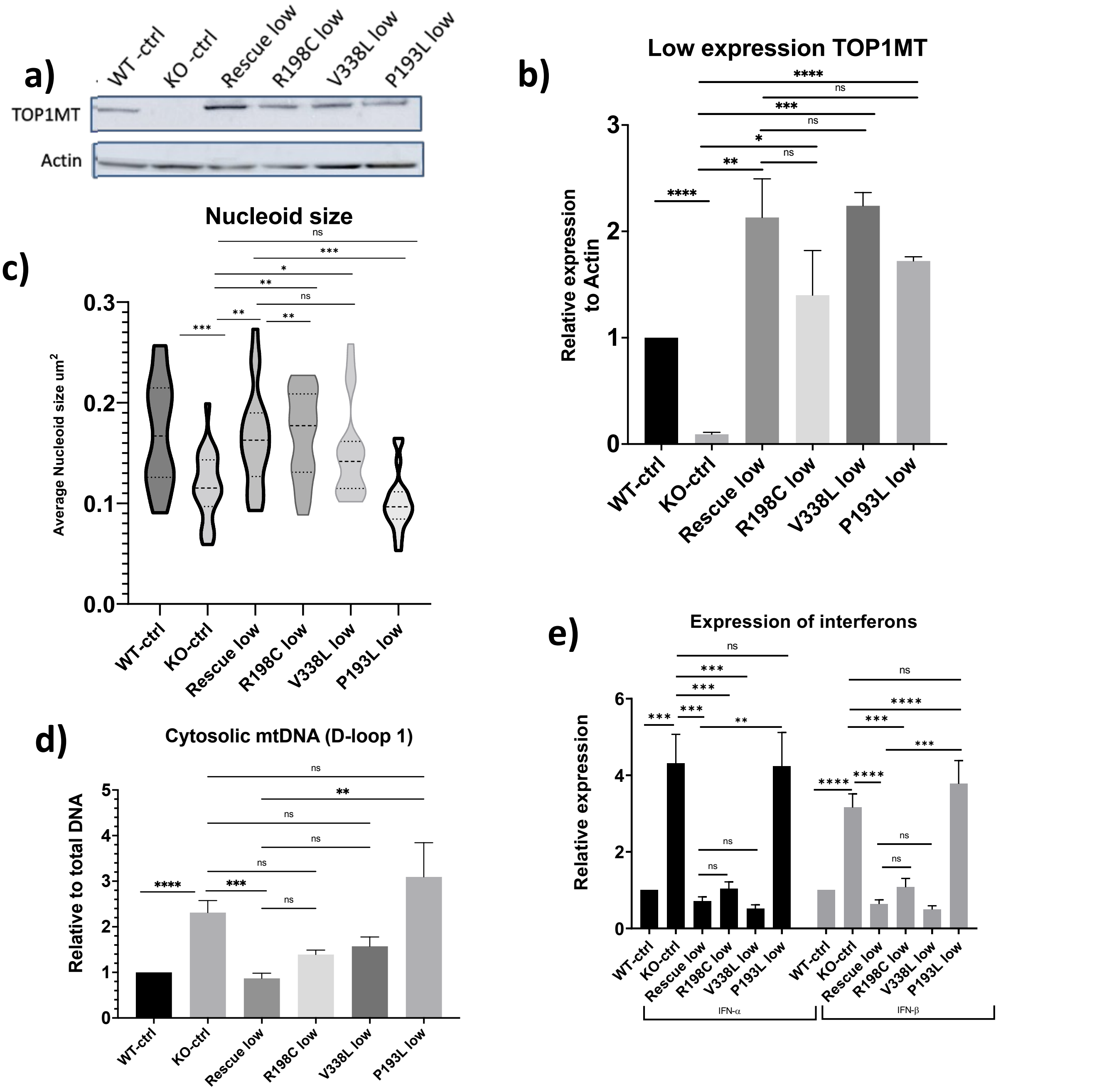


**Figure S6: Cytosolic mtDNA release and interferon levels in cell lines expressing low levels of TOP1MT.** a) Representative western blots showing reduced levels of the TOP1MT protein in cells sorted for lower expression of WT, R198C, V338L and P193L variants. Uncropped images of western blots are available in Fig S8d. b) Quantification of western blots of TOP1MT protein as in (a), from three independent experiments, corrected to Actin as a load control. c) Violin plots showing quantification of the average mtDNA nucleoid size from 25 cells stained with PicoGreen (dsDNA: nuclear and mtDNA). d) qPCR quantification of cytosolic mtDNA levels from indicated cells using the D-loop region primer set, shows that only the P193L TOP1MT variant is unable to rescue the increased cytosolic mtDNA when expressed at low levels. Normalization was done relative to the corresponding amplicons in total DNA from the same cells. e) qRT-PCR analysis of type I interferon expression from indicated cells shows that only the P193L TOP1MT variant is unable to rescue the elevated type I interferon levels when expressed at low levels. All statistical analysis were done using unpaired student t-test. P-values *<0.05, ** <0.01, *** <0.001 and **** <0.0001. 'ns' signifies no significant differences between indicated groups. Error bars represent standard error of mean.

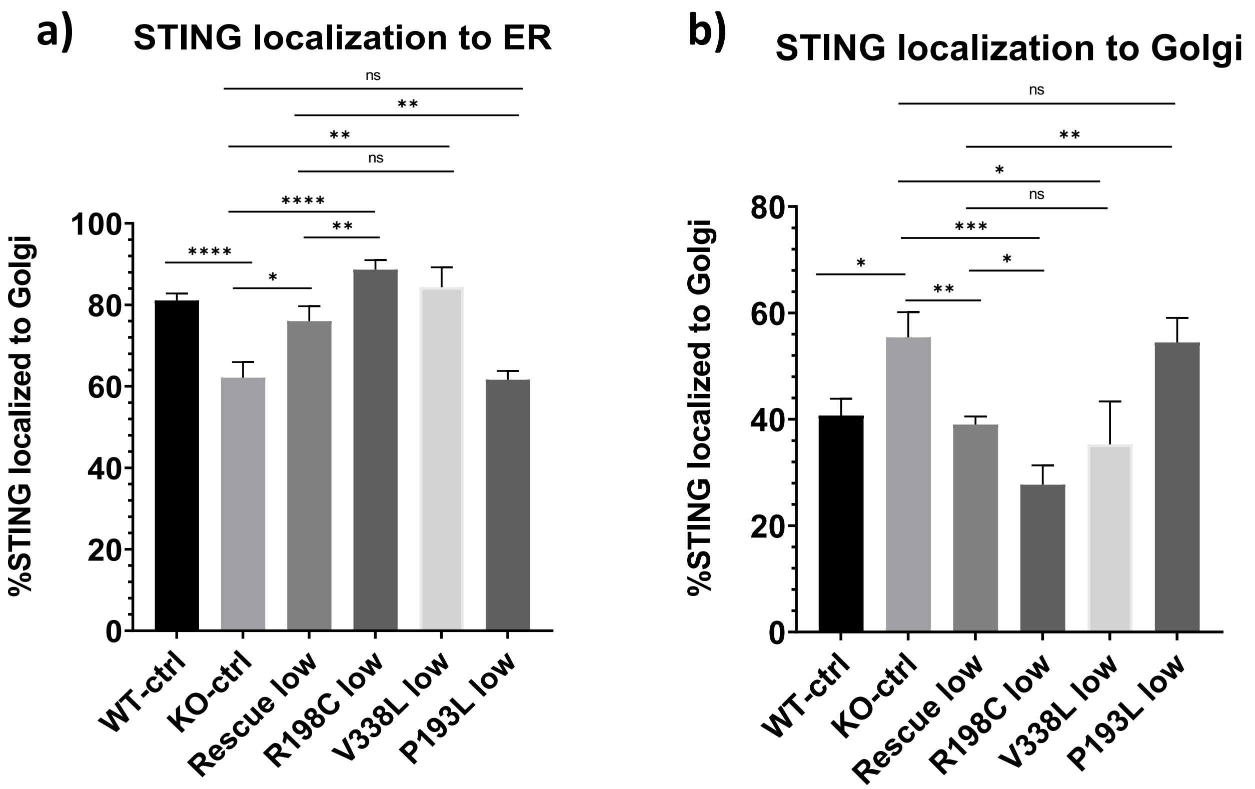


**Figure S7: STING relocates from the ER to the Golgi in TOP1MT KO and P193L cells but not in R198C and V338L cells. a,b)** Quantification of the percentage of STING localizing to the ER (a) or Golgi (b) from at least 10 images were fixed. Cells were immunofluorescent stained with anti-STING and anti-Calnexin (ER) or anti-TGN46 (Golgi) primary antibodies as in Fig 8 in the main text. While low expression of WT (Rescue), R198C or V338L TOP1MT variants restored STING localization, low expression of the P193L variant was unable to rescue the phenotype. Statistical analysis was done using unpaired student t-test and p values*<0.05, ** <0.01, ***<0.001 and **** <0.0001. ‘ns’ signifies no significant differences between indicated groups. Error bars represent standard error of mean.

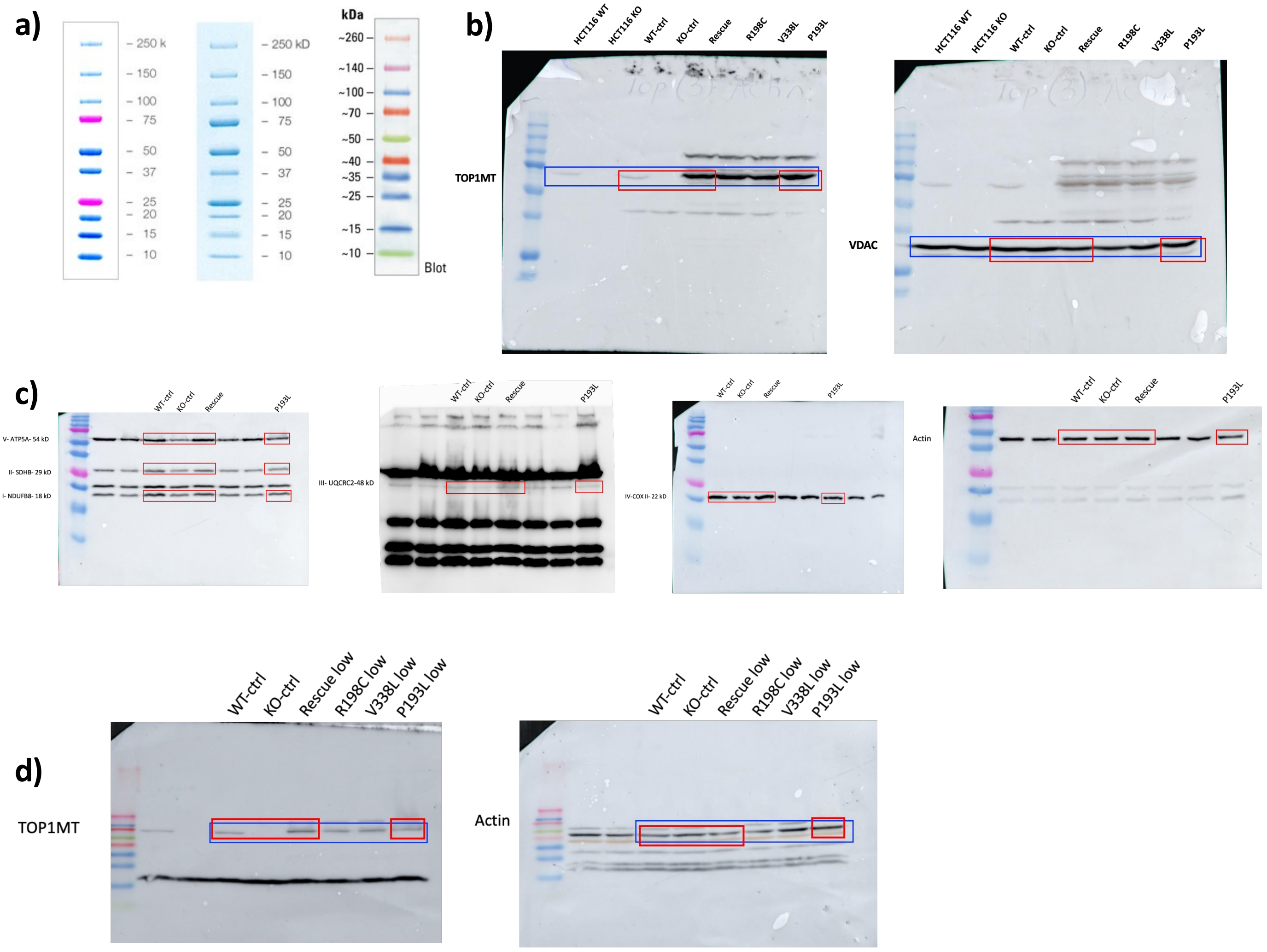


**Figure S8. Uncropped western blots. a)** Protein ladders of the various molecular weight markers used for size reference. **b)** Uncropped western blots for TOP1MT and VDAC used for figures 4c (red) and S5a (blue). **c)** Uncropped western blots for the indicated oxidative phosphorylation proteins and actin used for Fig 6d. **d)** Uncropped western blots for TOP1MT and Actin used in figures 7a (red) and S6a (blue).
