## Supplemental Information for "Activation of the cGAS-STING innate immune response in cells with deficient mitochondrial topoisomerase TOP1MT"

**Supplementary Information File 1**

The following clinical summaries have been prepared by WTG, in consultation with the family. WTG takes responsibility for the accuracy of the information presented herein. All study subjects have provided informed consent for research analysis and publication, in accordance with Declaration of Helsinki principles.

**(III-7) (Father):** III-7 is generally in good health. He is 56 years of age, 183 cm tall and weighs 91 kg (self-declared recent weight). He was hospitalized for 2 weeks as an inpatient with COVID-19 during the recent pandemic. He is currently taking no medications, vitamins or supplements. In addition to being a heterozygous carrier of the TOP1MT variant described in this paper, III-7 is a heterozygous carrier of congenital leptin deficiency (data on this research participant has previously been presented in[1], and in[2].

**(III-8) A1020 (Mother):** A1020 is generally in poor health. She is 56 years of age, 175 cm tall and weighs 117 kg (self-declared recent weight). Like her husband, she was hospitalized for 2 weeks as an inpatient with COVID-19 during the recent pandemic. She has hypothyroidism and impaired fasting glucose (6.5 mmol/L) with haemoglobin A1c of 5.4%. She suffers from anxiety and depression, and has been diagnosed with systemic lupus erythematosus, Sjogren’s syndrome, fibromyalgia, celiac disease and moderate to severe interstitial lung disease (pulmonary fibrosis). She has required home oxygen via nasal prongs since August of 2019, prior to the COVID-19 pandemic. Medications include mycophenolate mofetil, dapsone, gabapentin, amitryptiline, zopiclone, duloxetine, warfarin and Synthroid. She also takes vitamin D and biotin supplements, and follows a gluten-free diet. In addition to being a heterozygous carrier of the TOP1MT variant described in this paper, A1020 is a heterozygous carrier of congenital leptin deficiency (data on this research participant has previously been presented in[1], and in[2].

**(IV-3) (Eldest Sister):** IV-3 is generally in good health and 28 years of age. She was diagnosed with homozygosity for congenital leptin deficiency at age 3 years, and has been treated with recombinant human leptin since age 4 years (data on this research participant has previously been presented in[2]. In addition to congenital leptin deficiency, she was diagnosed with rheumatoid arthritis at age 5 years, as well as celiac disease. She is currently taking recombinant human leptin, certolizumab and naproxen. In addition to being homozygous for congenital leptin deficiency, she is homozygous for the TOP1MT variant described in this paper.

**(IV-4) (Middle Sister):** IV-4 is generally in good health and 24 years of age. She was diagnosed with rheumatoid arthritis at age 9 years, as well as celiac disease. She is prone to anxiety and depression. She follows a gluten-free diet, takes monthly infusions of tocilizumab, and takes antidepressant medications when needed. She has not yet sought clinical-grade testing for the leptin deficiency variant known to be present in her family, but is known not to be homozygous for that variant based on her phenotype. She is heterozygous for the TOP1MT variant described in this paper.

**(IV-5) (Youngest Sister):** IV-4 is generally in good health and 17 years of age. She was diagnosed with rheumatoid arthritis at age 2 years, and type 1 diabetes at age 5.5. years. She takes insulin therapy as well as monthly infusions of tocilizumab. She has not yet sought clinical-grade testing for the leptin deficiency variant known to be present in her family, but is known not to be homozygous for that variant based on her phenotype. She is homozygous for the TOP1MT variant described in this paper.

**References:**

1.	Farooqi, I.S., et al., *Partial leptin deficiency and human adiposity.* Nature, 2001. **414**(6859): p. 34-5.

2.	Gibson, W.T., et al., *Congenital leptin deficiency due to homozygosity for the Delta133G mutation: report of another case and evaluation of response to four years of leptin therapy.* J Clin Endocrinol Metab, 2004. **89**(10): p. 4821-6.
